# Mutation-Induced Pocket Deactivation: How Ser353/Pro245 Alters K_Ca_2.2 vs K_Ca_3.1 Ligand Selectivity

**DOI:** 10.64898/2026.05.03.722491

**Authors:** Matteo Gozzi, Joana Massa, Oliver Koch

**Affiliations:** Chemical biology of ion channels (Chembion), GRK 2515, University of Münster, Münster, Germany; Institute of Pharmaceutical and Medicinal Chemistry, University of Münster, Münster, Germany

**Keywords:** K_Ca_2.2, K_Ca_3.1, selectivity, molecular dynamics simulations, docking, in silico mutagenesis

## Abstract

The K_Ca_2.2 and K_Ca_3.1 channels are fundamental regulator of cellular K^+^ concentration, and promising target to treat diseases such as spinocerebellar ataxia and cancer. To fully exploit their therapeutic potential, and to continue studying their pathophysiological role, it is crucial to develop selective modulators for each of these two channels. Here we present a computational study to identify the molecular determinants behind the selectivity of two recently reported K_Ca_2.2 modulators. We leveraged a protocol combining *in silico* mutagenesis, molecular dynamics simulations, and protein-ligand docking to analyse the pockets targeted by these ligands. We identified a Ser353/Pro245 substitution to be the main driver of the distinct pocket shapes in K_Ca_2.2 and K_Ca_3.1 channels, ultimately defining modulator selectivity. This approach provides novel insights into the structural differences of this binding site across potassium channel subtypes, shedding light on the selectivity determinants of modulators targeting this pocket.

## 1 INTRODUCTION

Potassium (K^+^) selective ion channels are membrane proteins responsible for the regulation of cellular K^+^ concentration. Through this role, they control fundamental physiological functions, including hormone secretion, cellular excitability, epithelial transport, and cell volume variation^[1]^. Among these proteins are the small-conductance calcium-activated potassium channels K_Ca_2.1, K_Ca_2.2, and K_Ca_2.3 (also named SK1-3, *KCNN1-3*), and the intermediate-conductance calcium activated potassium channel K_Ca_3.1 (also named IK, SK4, *KCNN4*)^[1]^. Their K^+^ conductance is modified in response to changes in cytoplasmatic Ca^2+^ concentration, establishing a tight coupling between the concentration of these two ions^[1,2]^.

K_Ca_2.1-3 and K_Ca_3.1 channels are characterised by a homotetrameric structure, in which each subunit is composed of six transmembrane helices (S1-S6), and three additional intracellular helices (HA, HB, HC). S5 and S6 of every subunit assemble to form the ion-conducting pore, while the selectivity filter region located in the S5-S6 linker is responsible for K^+^ specificity^[3]^. Calcium sensitivity is mediated by the accessory protein calmodulin (CaM) which, through its C-terminal lobe (C-Lobe), constitutively binds the HA and HB helices of each channel subunit^[4]^.

The first experimental structure of a human calcium-activated potassium channel was obtained for the K_Ca_3.1 channel^[5]^, and recently the first human K_Ca_2.2 structure was resolved^[6]^ (**Figure 1A**). These structures enabled the elucidation of the gating mechanism of these channels. Upon calcium binding to CaM, the N-terminal lobe (N-Lobe) of this subunit forms a stable interaction with the channel S4-S5 linker. This induces a reciprocal distancing of the S6 helix, on which the hydrophobic gate is located (residue Val390 in K_Ca_2.2 and Val282 in K_Ca_3.1). Widening of this constriction site, induces an open conformation of the pore, allowing for K^+^ flux^[5,6]^. Recently, molecular dynamics (MD) simulations elucidated atomistic details of the K^+^ pathway. The simulations suggested that K^+^ ions enter the channel through two lateral intracellular fenestrations, cross the hydrophobic gate and ultimately exit the channel through the selectivity filter^[7]^.

**FIGURE 1.**
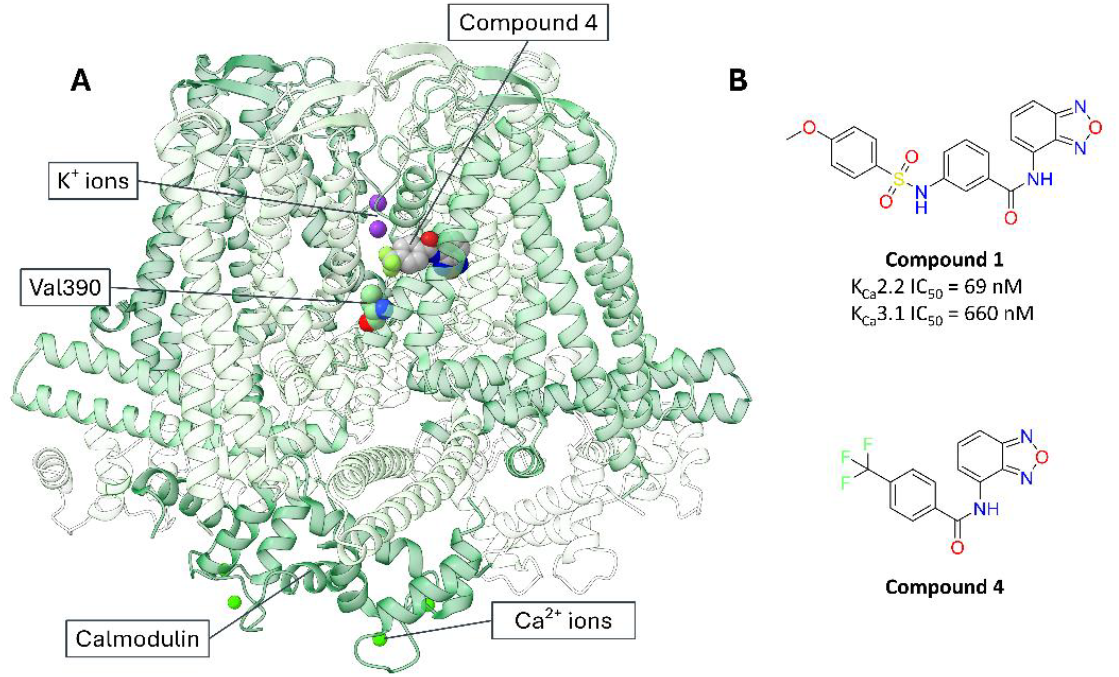
**A**)Structure of the Compound 4-bound K_Ca_2.2 channel (green ribbons, PDB ID: 9O5O^[6]^). Co-determined K_+_ ions located in the ion selectivity filter, Compound 4, gating residue Val390 and co-determined Ca_2+_ ions are represented with Van der Waals spheres. For better clarity, the represented Compound 4, residue Val390 and Ca_2+_ ions correspond to one subunit only. **B**) Chemical structure of K_Ca_2.2 inhibitor Compound 1 and K_Ca_2.2 activator Compound 4^[6]^

Over the decades, numerous studies have demonstrated the therapeutic potential of K_Ca_2.2 and K_Ca_3.1 modulation. K_Ca_2.2 inhibitors have shown potential against atrial fibrillation^[8]^ and Alzheimer’s disease^[9,10]^, with the molecule AP30663 successfully completing phase II clinical trials^[11]^. Meanwhile, K_Ca_2.2 activators could be employed to treat spinocerebellar ataxia^[12]^, and alcohol dependence^[13]^, or serve as neuroprotectants in stroke episodes^[14]^. Inhibition of K_Ca_3.1 channels has produced promising results in sickle-cell anaemia^[15–17]^, and also in cancer treatment^[18,19]^, with the blocker senicapoc approaching clinical trials for glioblastoma in combination with perampanel^[20]^. Additionally, K_Ca_3.1 activators showed potential in treating hypertension^[21,22]^ and cystic fibrosis^[23,24]^.

The diverse pathological implications of the K_Ca_2.2 and K_Ca_3.1 channels require the development of selective modulators to harness their full therapeutical potential, and to further study their function. Indeed, K_Ca_2.2 activators could be beneficial in CNS-involving diseases, but a lack of selectivity towards peripheral K_Ca_3.1 channels might lead to unwanted vascular side effects such as hypotension^[22,25]^. Conversely, inhibiting K_Ca_3.1 to treat glioblastoma could lead to seizures and neurodegeneration if K_Ca_2.2 channels are also blocked^[26]^.

To understand the determinants underlying the selectivity of K_Ca_2.2 and K_Ca_3.1 modulators, diverse techniques have been applied. Thanks to recent cryo-EM structures, the interaction between specific positive allosteric modulators (PAM) and their respective targets K_Ca_2.2^[27]^ and K_Ca_3.1^[28]^ has been elucidated, providing a rationale for ligand selectivity. In a study by Nguyen et al., homology modelling and molecular docking were successfully combined to predict the binding mode of a set of K_Ca_3.1 pore blockers, validating the computational results with site-directed mutagenesis (SDM) experiments^[29]^.

Recently, the cryo-EM structures of the K_Ca_2.2 channel in complex with a novel small-molecule inhibitor (Compound 1) and an activator (Compound 4) derived from a high-throughput screening campaign were reported^[6]^ (**Figure 1A,B**). Compound 1 inhibits K_Ca_2.2 with an IC_50_ of 69 nM, and crucially, showed a 10-fold selectivity towards K_Ca_3.1 (IC_50_ of 660 nM). Compound 4 is only described as an activator without specified EC_50_ values. Notably, both molecules bind to a previously undescribed region of the K_Ca_2.2 channel, comprised between the S5, P-Helix and S6 helices (S5-Phelix-S6 pocket). Both molecules are characterised by a common scaffold, in which a benzoxadiazole moiety is linked to a phenyl ring through an amide bond. This core offers the complementarity needed for interacting with the surrounding residues Leu321, Ala325, Ile352, Phe356, and Thr378, establishing also a π-π interaction with residue Trp322. The opposite activity of the two ligands depends on the diverse substituents located on the phenyl ring, which interact with specific residues located on the S6 helix. The meta-methoxybenzene-1-sulfonamide moiety characterising Compound 1, which establishes an additional hydrogen bond (H-bond) with Ser318 and a π-π interaction with Phe356, is capable of forcing the closure of K_Ca_2.2 hydrophobic gate by interacting with Thr386. On the contrary, the trifluoro-methyl group of Compound 4 interacts with Ile380 of a neighbouring subunit, stabilising the open state of the gate^[6]^.

Starting from these newly reported K_Ca_2.2 modulators, we initially sought to determine whether the S5-Phelix-S6 pocket could be leveraged to develop novel K_Ca_3.1 modulators. Additionally, we wanted to gain a deeper insight into the molecular basis behind the selectivity of the K_Ca_2.2 targeting compounds. Thanks to a combined computational approach, we were able to highlight key differences between K_Ca_2.2 and K_Ca_3.1 S5-Phelix-S6 pockets, and to identify specific residues potentially responsible for modulator selectivity.

## 2 RESULTS AND DISCUSSION

### 2.1 S5-Phelix-S6 Pocket Comparison Between K_Ca_2.2 and K_Ca_3.1

As a first step, the molecular basis underlying the selectivity of Compound 1 against K_Ca_3.1 was investigated. Therefore, the residues lining the S5-Phelix-S6 binding pocket in K_Ca_2.2 were compared to the corresponding positions in K_Ca_3.1. In the following sections, residues differences between the two channels will be mentioned using the three-letter code, indicating first the K_Ca_2.2 residue, followed by the K_Ca_3.1 one (e.g. Trp322/Trp216). Using the Compound 1-bound K_Ca_2.2 structure, and the K_Ca_3.1_closed structure, 23 residues within 5 Å from Compound 1 were identified (**Figure 2D**). The two groups of residues share a high sequence similarity (78% considering BLOSUM62^[30]^ score > 0), with only nine sequence substitutions between the two channels out of the 23 amino acids lining the pocket (**Figure 2A,D**). Nonetheless, the two pockets are characterised by important structural differences. Most notably, Trp322/Trp216, which is part of S5, is conserved in both channels but assumes two distinct conformations (**Figure 2B**). In the K_Ca_2.2 channel the side chain is directed towards the extracellular side, allowing Compound 1 and 4 to easily bind. In contrast, in K_Ca_3.1 the indole ring is directed towards the intracellular side and would clash with both ligands as it obstructs the S5-Phelix-S6 pocket (**Figure 2B**). This potentially determines the selectivity of Compound 1, as also hypothesised by Cassell et al.^[6]^.

**FIGURE 2.**
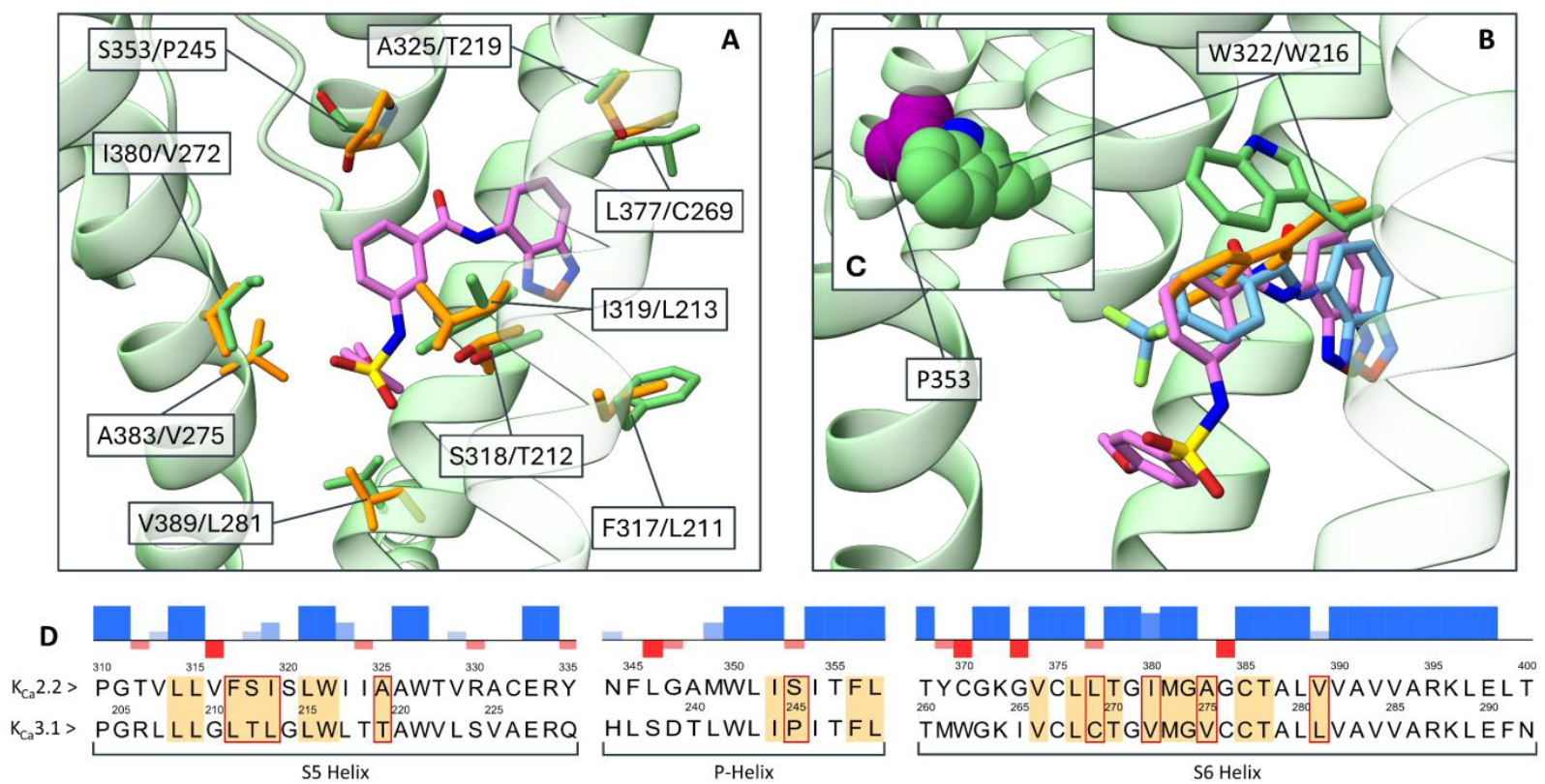
S5-Phelix-S6 pocket in K_Ca_2.2 (green ribbons and sticks, PDB ID: 9O53^[6]^) and K_Ca_3.1 (orange sticks, PDB ID: 6CNM^[5]^). Compound 1 is represented with pink sticks, while Compound 4 with light blue sticks. Residues are indicated using the one-letter core, with first residues from K_Ca_2.2 followed by the ones from K_Ca_3.1. **A**) Amino acid substitutions between the K_Ca_2.2 and K_Ca_3.1 channels. **B**) Clash between residue Trp216 in K_Ca_3.1 and Compound 1/Compound 4; **C**) Focus on the clash between Trp322 (green Van der Waals spheres) and the mutated Ser353Pro residue (purple Van der Waals spheres) in structure K_Ca_2.2. **D**) Alignment of K_Ca_2.2 and K_Ca_3.1 channel sequences. Residues lining the S5-Phelix-S6 pocket (within 5 Å from Compound 1) are highlighted in orange. Residues substitution between K_Ca_2.2 and K_Ca_3.1 are indicated with a red box. The sequence alignment was computed using the EMBL-EBI Clustal Omega web server^[33]^.

Although experimental selectivity data are available only for the inhibitor Compound 1, the common binding mode of the two ligands suggests that Compound 4 shares a similar selectivity profile. Interestingly, the Trp322/Trp216 conformation does not depend on the channel state, as it is consistent across all available wild-type K_Ca_2.2 and K_Ca_3.1 experimental structures (**Table S1**,**2**).

Further differences were identified when comparing the binding sites in K_Ca_2.2 and K_Ca_3.1 (**Figure 2A,D**). We hypothesised that these have only a minor impact on the binding of Compound 1 and Compound 4. Specifically, the Ile319/Leu213 and Ile380/Val272 substitutions could potentially modify the shape complementarity between the pocket and the phenyl moiety of the two ligands, while Ala383/Val275 and Val389/Leu281 might influence the placement of the methoxy-phenyl moiety of Compound 1. The Ser318/Thr212 substitution might impact ligand binding due to the additional methyl group of Thr212, as already suggested by Cassel et al.^[6]^. However, the methyl group faces away from the ligand site in all available K_Ca_3.1 structure. Lastly, since residues involved in the Phe317/Leu211 and Leu377/Cys269 substitutions project their side chains away from the ligand binding site, it is unlikely that they have an impact on ligand selectivity.

Apart from these substitutions, the K_Ca_2.2 residue Ser353 is replaced by Pro245 in K_Ca_3.1 (**Figure 2A,D**). Trp322/Trp216 are in close proximity to Ser353/Pro245, which hints at a major contribution of these last residues to the diverse tryptophan conformations. Upon *in silico* mutation of Ser353 to proline in the Compound 1-bound K_Ca_2.2 structure, major clashes with Trp322 were identified (, **Figure 2C**). This result suggests that the presence of a proline residue in this specific position is not compatible with the Trp322 conformation captured in the K_Ca_2.2 structures, and thus Pro245 could be the involved in the downward-facing conformation of Trp216 in K_Ca_3.1.

Interestingly, this Ser/Pro substitution between K_Ca_2.2 and K_Ca_3.1 is conserved across species (**Figure S1**), hinting at a potential role in steering differences between these channels. By combining computational techniques and SDM, Garneau et al. demonstrated that the aromatic interaction between Trp216 and Phe248 in K_Ca_3.1 plays a crucial role in shaping the channel maximum open probability (POMAX)^[31]^. Thanks to the Pro353/Ser245 substitution, in K_Ca_2.2 Trp322 can adopt a different conformation compared to K_Ca_3.1 and loses this aromatic interaction, which is in line with the higher POMAX of this channel (0.8 in K_Ca_2.2^[32]^ vs 0.1-0.2 in K_Ca_3.1^[31,32]^). This structural comparison of the S5-Phelix-S6 pockets between the K_Ca_2.2 and K_Ca_3.1 channels highlights major differences between the two binding site. Since residue Trp216 side chain obstructs this pocket in the K_Ca_3.1 structures, we hypothesise that the selectivity profiles of Compounds 1 and 4 are directly deriving from the diverse Trp322/Trp216 conformations between the K_Ca_2.2 and K_Ca_3.1 channels, which in turn could be induced by the crucial Ser353/Pro245 substitution.

### 2.2 In Silico Mutagenesis and Molecular Dynamics Analysis

To probe the role of Pro245 in restraining the Trp216 conformation in K_Ca_3.1, we leveraged a combination of *in silico* mutagenesis and MD simulations. A closed and an open conformation of the K_Ca_3.1 channel (described in the *Methods* section) were chosen for this analysis. This enables a direct comparison to the closed and open Compound 1-bound and Compound 4-bound K_Ca_2.2 structures. The Pro245Ser mutation was applied *in silico t*o the K_Ca_3.1 structures, and the wild-type (WT) and mutated (P245S) structures were subjected to triplicate MD simulations with 500 ns long production phases. In the following sections the four systems will be referred to as K_Ca_3.1_closed_WT, K_Ca_3.1_closed_P245S, K_Ca_3.1_open_WT, and K_Ca_3.1_open_P245S, respectively.

#### 2.2.1 Protein stability and folding in the WT and P245S systems

Analysis of the root mean square deviation (RMSD) profiles highlighted the stability of the channel subunits in both the WT and the P245S systems, with the RMSD value converging to ∼4 Å after approximately 150 ns (**Figure S2**,**3**), indicating that the Pro245Ser mutation did not induce significant perturbation of the channel stability. The calmodulin subunits exhibit less stable RMSD profiles in the K_Ca_3.1_closed simulations compared to the K_Ca_3.1_open ones, and are also characterised by higher RMSD average values compared to the channel subunits (**Table S3**). This behaviour is consistent with the non-calcium-bound state of CaM in the K_Ca_3.1_closed systems, which is known to lead to increased instability of this subunit^[5]^. To evaluate the impact of the Pro245Ser mutation on protein folding, the radius of gyration (RoG) of the K_Ca_3.1 channel was analysed throughout both the WT and P245S simulations, and no major differences between two sets were noticed (**Figure S4**). Altogether, this data suggest that the Pro245Ser mutation does not induce instability or unfolding of K_Ca_3.1 channel compared to WT.

#### 2.2.2 Trp216 Conformational change

In order to study the conformational state of Trp216 between the WT and P245S systems, the trajectories were visually inspected, followed by the analysis of the Trp216 *χ*_1_ and *χ*_2_ dihedral angles. In every replicate of the WT systems, the Trp216 rotamers cluster around the conformation captured in the experimental K_Ca_3.1 structures, named *conformation a* (**Figure 3A,C**), which is incompatible with ligand binding. Although in the K_Ca_3.1_closed system the Pro245Ser mutation did not induce substantial changes in the Trp216 conformer (**Figure 3A,B**), in the K_Ca_3.1_open system major differences can be observed between the WT and P245S simulations (**Figure 3C,D**). While Trp216 maintains the initial *conformation a* (**Figure 3C and 4A**) in the K_Ca_3.1_open_WT simulations, a conformational switch was recorded in one of the K_Ca_3.1_open_P245S replicates. Specifically, Trp216 deviates from the initial *conformation a* after approximately 10 ns, alternating between two rotamers, named *conformation b* and *c* (**Figure 3D and 4B**,**C**), which project their side chains towards the extracellular side. These new conformations show high similarity to the one observed in the K_Ca_2.2 structures. Additional MD simulation of Compound 1 and 4-bound K_Ca_2.2 revealed comparable Trp322/Trp216 dihedral plots between these systems and K_Ca_3.1_open_P245S (**Figure S5**). These results indicate a different Trp216 behaviour between the WT and P245S systems. The low sampling of the rotamer transition in only one out of the three P245S replicates suggests that this flip represents a low probability event in K_Ca_3.1, although the upward-oriented conformation itself is stable in the K_Ca_3.1_open_P245S simulation. On the other hand, in K_Ca_2.2 this conformation seems to be the default rotamer state as it is observed in all experimental structures of this channel (**Table S1**). Ultimately, the simulations suggest that in presence of the Pro245Ser mutation Trp216 is capable of spontaneously changing rotamer, reinforcing the hypothesis that the Ser353/Pro245 substitution between K_Ca_2.2 and K_Ca_3.1 plays a major role in the S5-Phelix-S6 pocket structure.

**FIGURE 3.**
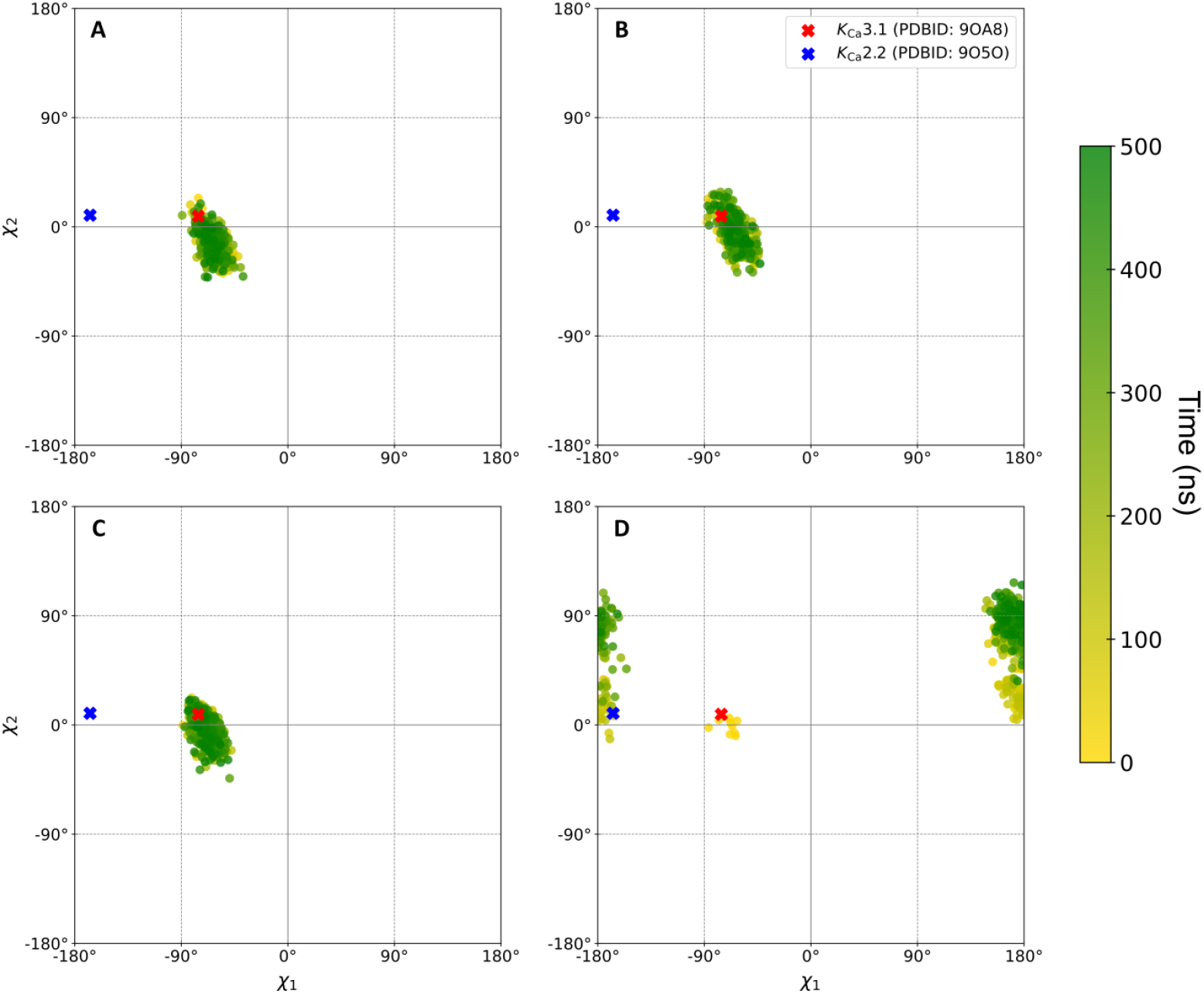
Plots of the *χ*_1_ and *χ*_2_ dihedral angles of residue Trp216 during the K_Ca_3.1 MD simulations. Each dot represents a combination of dihedral angles in a specific frame of the trajectory. Time evolution along the simulation is represented by colour shift from yellow to green. Reference dihedral angles obtained from the Compound 4-bound K_Ca_2.2 and K_Ca_3.1_open cryo-EM structures are indicated with a blue and red cross, respectively. **A**) System K_Ca_3.1_closed_WT, run 1, chain A. **B**) System K_Ca_3.1_open_P245S, run 1, chain A. **C**) System K_Ca_3.1_open_WT, run 1, chain A. **D**) System K_Ca_3.1_open_P245S, run 2, chain B.

#### 2.2.3 Trp216 interactions analysis

To gain a better understanding of the mechanism behind the Trp216 conformational switch, the interactions established by this residue during the K_Ca_3.1_open_WT and K_Ca_3.1_open_P245S simulations were analysed and compared.

H-bonds between the backbone of Trp216 and the backbones of Thr212 and Ala220 are conserved in both systems, as they contribute to maintaining the α-helical structure of S5 (**Figure 4D,E**). In the WT simulations, the *conformation a* of Trp216 is stabilised by extensive hydrophobic contacts with Leu213 (located on the S5 helix), Phe248, Leu249 (both located in the P-Helix), and Leu268 (located in the S6 helix of a neighbouring subunit), with an additional T-shaped π-π interaction established with Phe248 (**Figure 4A,D**). Analysis of the interaction pattern in the K_Ca_3.1_open_P245S system reveals major differences compared to the K_Ca_3.1_open_WT. First, the hydrophobic interactions with Leu249 and the T-shaped π-π interaction with Phe248 are lost after ∼10 ns, due to the rotation of Trp216 side chain (**Figure 4E**). By analysing the distance between Ser245 (located on the P-Helix) and Trp216 during the K_Ca_3.1_open_P245S trajectory, it is possible to observe that the distance is stable at ∼6.5 Å in every chain except for chain B, were Trp216 conformational shift occurs. Here the distance rises to ∼9.5 Å, allowing for the upward rotation of Trp216 side chain, and then stabilises at ∼7.5 Å due to the new Trp216 conformation (**Figure S6**).

**FIGURE 4.**
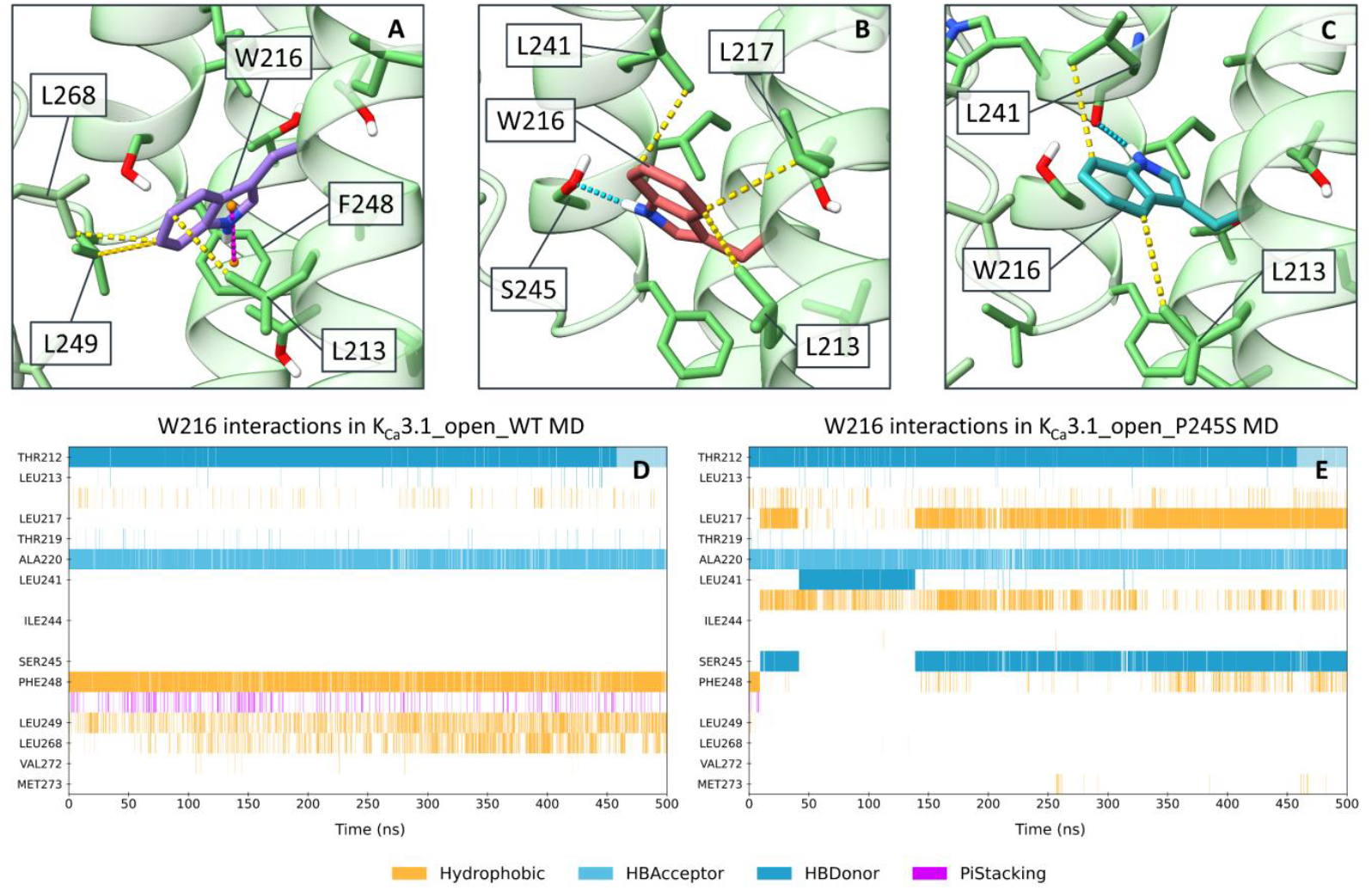
Interactions established by residue Trp216 during the K_Ca_3.1_open MD simulations. 3D representation of the interactions established by Trp216 side chain in **A**) *conformation a*, **B**) *conformation b*, and **C**) *conformation c*. The K_Ca_3.1 channel is represented by green ribbons, with residues in green sticks. The different Trp216 conformations are highlighted in purple, orange and cyan, respectively. Bar plots representing the time evolution of the interactions established by Trp216 across **D**) the K_Ca_3.1_open_WT simulations (run1, chain A), and **E**) the K_Ca_3.1_open_P245S simulations (run1, chain B). Non-covalent interactions are indicated with dashed lines, specifically H-bonds are represented in cyan, Hydrophobic in yellow and π-π stacking in magenta.

After this transition from the initial *conformation a*, first an H-bond with Ser245 side chain oxygen and later with Leu241 backbone carbonyl oxygen are detected. These two polar interactions stabilise the newly adopted Trp216 *conformations b* (**Figure 4B**) and *c* (**Figure 4C**). The last of these two interactions corresponds to the state captured in the Compound 1 and 4-bound K_Ca_2.2 structures, where a H-bond between the Trp322 side chain and Met349 (corresponding to Trp216 and Leu241 in K_Ca_3.1) is detected. Furthermore, the upward-oriented Trp216 *conformations b* and *c* establish stable hydrophobic interactions with Leu241 (**Figure 4B,C,E**), which is replaced by Met349 in K_Ca_2.2. This substitution might result in higher stabilization of the upward-oriented Trp322 conformation in the K_Ca_2.2 channel, since Met349 can establish sulphur-π interactions with this residue, which are stronger compared to the aliphatic-aromatic interactions mediated by Leu241 in K_Ca_3.1^[34]^.

Taken together, this analysis indicates that i) the initial Trp216 *conformation a* is stabilised by a set of hydrophobic and aromatic interactions with residues P248, Leu249 and Leu268, and ii) adoption of other rotamers (*conformations b and c*) is impaired by the short distance to the P-Helix. This explains the low probability of the observed Trp216 conformational switch. However, upon an increase in the Trp216 - P-Helix distance of ∼2.5 Å and in the presence of the Pro2345Ser mutation, the side chain of Trp216 is capable of adopting new rotamers, stabilised by two energetically favourable H-bond interaction.

### 2.3 Analysis of the S5-Phelix-S6 Pocket in the K_Ca_3.1_open_P245S system

To investigate if the conformational changes observed in the MD simulations of the K_Ca_3.1_open_P245S system produced a site capable of binding Compound 1 and 4, the S5-Phelix-S6 pocket in was analysed in detail. First, volume and shape of the pocket throughout the trajectory were studied (**Figure 5A**). Starting from a value of 396 Å^3^, the pocket volume increases in the first 80 ns of the simulation, followed by a slight decrease and stabilisation around 700 Å^3^ in the second half of the trajectory. The maximum pocket volume achieved is 1240 Å^3^, which is comparable to the average volume of the pocket in the Compound 4-bound K_Ca_2.2 channel (1403 Å^3^). The pocket shape and location changes throughout the simulation and adopts conformations similar to the one captured in K_Ca_2.2 cryo-EM structures (**Figure S7**) which might allow for ligand binding in K_Ca_3.1.

**FIGURE 5.**
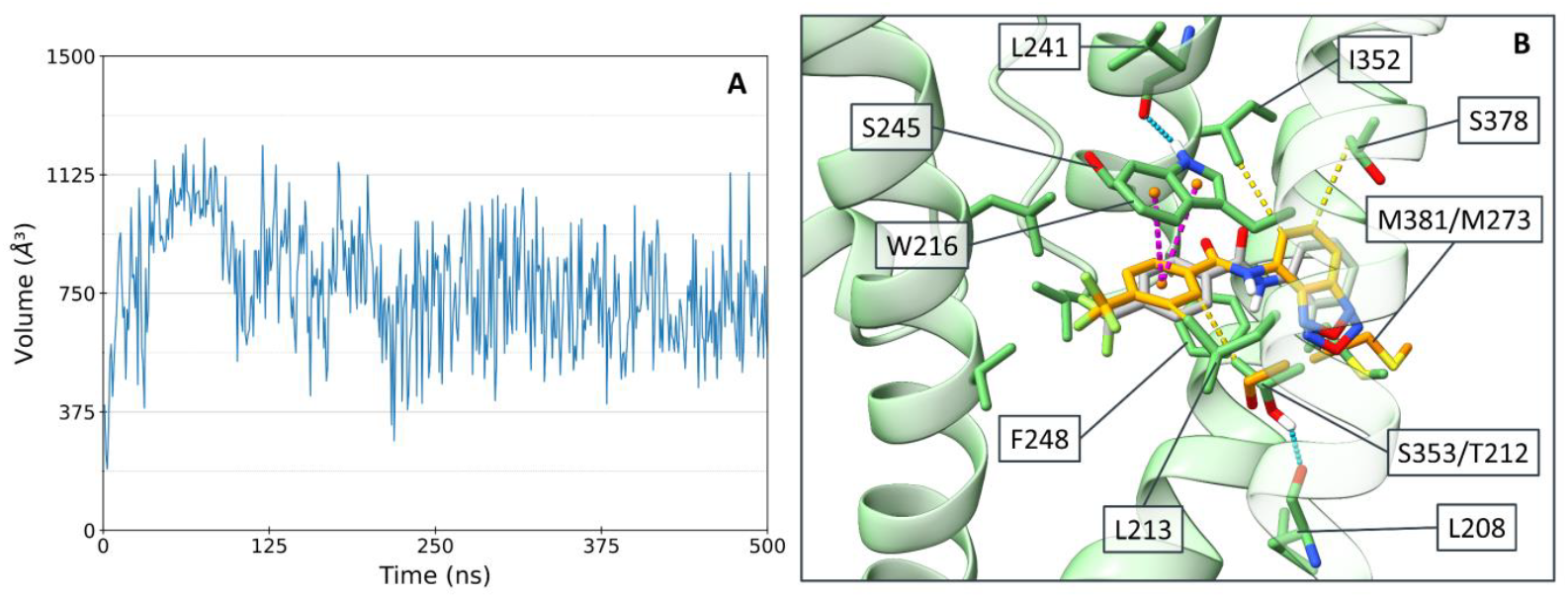
**A**) Volume variation of S5-Phelix-S6 binding pocket across the K_Ca_3.1_open_P245S simulation (run1, chain B). **B**) Comparison between the Compound 4 pose obtained via ensemble docking (gold sticks) and the reference cryo-EM binding mode (grey sticks). The K_Ca_3.1 channel is represented by green ribbons, with residues in green sticks. Non-covalent interactions are indicated with dashed lines, specifically H-bonds are represented in cyan, Hydrophobic in yellow and π-π stacking in magenta.

As a last step, ensemble docking was used to probe Compounds 4 binding at the K_Ca_3.1_open_P245S system. Ensemble docking is a variation of the classical docking technique in which, instead of using a single structure of the target protein to predict the binding mode of a ligand, multiple conformations of the same target are used. With this strategy it is possible to account for protein flexibility, and in this specific case to study the effects of the diverse Trp216 conformation on ligand binding. To obtain the structures for this calculation, frames from the K_Ca_3.1_open_P245S simulation were clustered based on the Trp216 rotamer. From each of the three resulting cluster a representative structure was extracted, which were then used for the ensemble docking of Compound 4.

Analysis of the docking results revealed that 80 out of the 100 generated poses depict the ligand in a binding mode closely resembling the pose in the Compound 4-bound K_Ca_2.2 channel cryo-EM structure (**Figure 5B**). Additionally, all the top-10 ranked poses are in line with the experimental binding mode, highlighting the ability of the scoring protocol to select conformations close to the experimental data. All generated binding modes are in the same channel snapshot (corresponding to 49 ns, *conformation c*), where the Trp216 projects the side chain towards the extracellular side, establishing a H-bond with Leu241 backbone carbonyl oxygen. Therefore, this channel state is suggested to be the most suitable for ligand binding. This result is in line with the fact that this Trp216 conformation most closely resembles the one captured by Compound 4-bound K_Ca_2.2. RMSD of the docking poses compared to the experimental binding mode ranges from 1.33 Å to 1.65 Å. Comparing the interactions established by Compound 4 in our docking poses and in the cryo-EM structure, only minor differences were identified. In both binding modes the benzoxadiazol moiety establishes extensive hydrophobic interactions with the surrounding residues (**Figure 5B**). The trifluoromethyl-benzyl moiety interacts via parallel π-π stacking with Trp216, although it lacks the same interaction with Phe248 (Phe356 in K_Ca_2.2), which is present in the cryo-EM structure. Minor differences in the placement could be attributed to the Ser353/Thr212 substitution between the K_Ca_2.2 and K_Ca_3.1, as already suggested in the “**S5-Phelix-S6 Pocket Comparison Between K**_**Ca**_**2.2 and K**_**Ca**_**3.1**” section. During the MD simulations, Thr212 establishes a stable H-bond with the backbone carbonyl oxygen of residue Leu208, rotating the methyl group toward the ligand binding site and thus reducing the available volume (**Figure 5B**).

Lastly, an additional docking round was conducted, employing the representative frame extracted from the Trp216 downward facing cluster of the MD simulation (corresponding to 2 ns, *conformation a*), and setting Trp216 as flexible. This did not lead to any ligand binding mode comparable to the one captured by the Compound 4-bound K_Ca_2.2 structure (**Figure S8**), indicating that in this conformation Trp216 is not capable of sampling rotamers compatible with the binding of Compound 4. Consequently, this highlights the essential role of MD simulations in sampling the effects of the Pro245Ser mutation, and in producing a channel conformation capable of accommodating Compound 4.

## 3 CONCLUSION

Developing selective modulators of K_Ca_2.2 and K_Ca_3.1 is crucial for exploiting their full therapeutic potential. In this work, we applied a combination of computational techniques to study the differences between the S5-Phelix-S6 pocket in K_Ca_2.2 and K_Ca_3.1, elucidating the molecular basis underlying the selectivity of two newly reported K_Ca_2.2 modulators, Compound 1 and Compound 4. Comparison of the pockets between the two channels identified two crucial differences, namely the distinct conformation of residues Trp322/Trp216 and the Ser353/P245 substitution. Due to the differences in Trp322/Trp216 rotamer, the full binding site is not available for ligand binding in K_Ca_3.1, potentially explain the lower potency of Compound 1 towards this channel. We hypothesised that the Ser353/Pro245 substitution in K_Ca_3.1 is responsible for restraining the conformation of Trp216. To challenge this hypothesis, residue Pro245 in the K_Ca_3.1 was mutated *in silico* to the corresponding serine residue found in the K_Ca_2.2 sequence. MD simulations of the WT and mutated channels suggested that the Pro245Ser mutation enables Trp216 to adopt conformations compatible with the one captured in the K_Ca_2.2 structures, although this conformational switch represents a low probability event. This low probability could be attributed to the close proximity between the S5 and P-Helix in K_Ca_3.1 cryo-EM structure, which hinders the free rotation of Trp216. This represents an intrinsic limitation of *in silico* mutagenesis, where proteins do not undergo folding, causing the WT and mutated structures to share the same backbone coordinates. Finally, we conducted ensemble docking of Compound 4 to conformations of the S5-Phelix-S6 pocket sampled from the mutated K_Ca_3.1 channel MD simulation, obtaining binding modes comparable to the ones captured in the cryo-EM structures. This suggests that the conformational shift of Trp216 enabled by Pro245Ser *in silico* mutation, allows the K_Ca_3.1 S5-Phelix-S6 pocket to transition from an “inactivated” to a “receptive” conformation capable of ligand binding.

To conclude, we propose the Ser353/Pro245 substitution between K_Ca_2.2 and K_Ca_3.1 channels to be the primary determinant of the selectivity of ligands binding to the S5-Phelix-S6 pocket, reinforcing the potential of targeting this previously undescribed site to develop molecules with selectivity for K_Ca_2.2 over K_Ca_3.1.

## 4 EXPERIMENTAL

### 4.1 Protein and Ligand Preparation

For this study, two structures of the K_Ca_3.1 channel and two of the K_Ca_2.2/K_Ca_3.1 chimera were used. The K_Ca_3.1 structures used represent the closed state (PDB ID: 6CNM^[5]^) and the open state (PDB ID: 9OA8^[27]^), which will be referred to as K_Ca_3.1_closed and K_Ca_3.1_open. The K_Ca_2.2/K_Ca_3.1 chimera structures used are the closed state with bound compound 1 (PBD ID: 9O53^[6]^), and the open state with bound compound 4 (PDB ID: 9O5O^[6]^), which will be referred to as Compound 1-bound K_Ca_2.2 and Compound 4-bound K_Ca_2.2.

After retrieving the structure from the Protein Data Bank (PDB) database^[35]^, lipids and glycosides were removed, proceeding with structure preparation using the MOE v2024.060^[36]^ suite. The missing S3-S4 loops (residues 124-141) in the K_Ca_3.1 structures, and the missing CaM loops (residues 113-118) in the K_Ca_2.2 structures were modelled using the “PDB search” function within “Loop Modeller”. Missing residue sidechains were modelled using “Structure Preparation”. Finally, protein and co-determined ligands were protonated using “Protonate3D” at pH 7.4. For the mutated K_Ca_3.1 systems, referred to as K_Ca_3.1_closed_P245S and K_Ca_3.1_open_P245S, MOE “Protein Builder” was used to mutate the residue Pro245 of the P-Helix to serine in every K_Ca_3.1 subunit.

Compound 4 was prepared for the docking calculation by extracting the ligand from the Compound 4-bound K_Ca_2.2 structure, followed by protonation using MOE “Protonate 3D” at pH 7.4, and energy minimisation of the conformer using the MMFF94x force field^[37]^.

### 4.2 Molecular Dynamics Simulations

Systems were prepared for simulations using CHARMM-GUI membrane builder^[38,39]^. Channel and calmodulin N-termini were acetylated (ACE) and C-termini methyl-amidated (CT3). In the K_Ca_2.2 systems a disulfide patch was applied between residues Cys320 and Cys302. Systems were oriented using PPM2.0^[40]^, based on the coordinates of the transmembrane domains (subunits A-D), and then inserted in a 1-palmitoyl-2-oleoyl-sn-glycero-3-phosphocholine (POPC) lipid bilayer. The protein-membrane systems were solvated with TIP3P^[41]^ water, adding K^+^ and Cl^−^ ions up to a concentration of 0.15 M to reach charge neutrality. The final simulation systems contained around 370,000 atoms the K_Ca_3.1 systems, and 310,000 for the K_Ca_2.2 systems. Additional details regarding the membrane systems are reported in **Table S4**.

Molecular dynamics (MD) simulations were conducted using the GROMACS v2024.4^[42]^ software. For the four K_Ca_3.1 simulations the Amber force field (FF) was applied, specifically ff14SB^[43]^ for proteins and Lipid21^[44]^ for lipids, while ligand parameters were calculated using Antechamber with AM1-BCC^[45]^ charges and GAFF^[46]^ atom types. For the two K_Ca_2.2 simulations CHARMM36m^[47]^ FF was used for protein and lipid molecules, while ligand parameters were calculated using CGenFF^[48]^.

Simulations were conducted on a local machine composed of two 18 cores Intel Xeon Gold 5220 CPUs and two RTX2080 Super GPUs, and on HPC nodes with two 64 cores AMD EPYC 7713 CPUs and eight Nvidia A40 GPUs, using 16 cores and one GPU each run. Each system was subjected to the default CHARMM-GUI minimization and equilibration protocol, followed by three independent 500 ns production runs in the NPT ensemble. In the production MD simulations, the temperature coupling was ensured with the V-rescale thermostat and held at 310.15 K with a 1 ps time constant. Semi-isotropic pressure coupling with the C-rescale barostat was used to keep the pressure at around 1 bar. A 5 ps time constant with a compressibility of 4.5 × 10^-5^ bar^-1^ was used. FF specific cutoffs for short-range Van der Waals and Coulomb interactions were used, corresponding to 9 Å in the K_Ca_3.1 simulations and 12 Å in the K_Ca_2.2 simulations. Long-range Coulomb interactions were handled with the Particle Mesh Edwald (PME)^[49]^ method. Bonds involving hydrogen atoms were restrained using the LINCS^[50]^.

### 4.3 Molecular Docking

Representative frames for the docking calculations were extracted from the MD simulations using the following approach. First, the Trp216 *χ*_1_ and *χ*_2_ dihedral angles obtained by the MD were decomposed in the respective sine and cosine components, which were then used for clustering the frames using the k-means algorithm^[51]^. The number of clusters was set to three. Next, the closest frames to the three cluster centroids (corresponding to 2 ns, 49 ns and 237 ns) were extracted from the MD trajectory. From the extracted protein structures only chains B and C were retained, since they compose the S5-PHelix-S6 pocket in which the Trp216 conformational switch occurs, and were used for the subsequent analysis steps.

Docking calculations were performed using GOLD, a genetic algorithm-based software^[52,53]^. The three representative frames extracted from the K_Ca_3.1_open_P245S MD were superimposed onto the 9O5O cryo-EM structure, and residues within 6 Å from the experimentally determined ligand were used to define the binding pocket. To allow for the correct ligand flexibility, amide bonds were allowed to flip. Ensemble docking was conducted using a genetic algorithm search efficiency of 200%, and early termination was disabled to allow for a more accurate conformational sampling. The number of individual genetic algorithm runs was set to 100, generating an equal amount of output binding modes, which were first scored using the ChemPLP scoring function^[54]^, followed by rescoring with GoldScore^[55]^. Compound 4 coordinates were extracted from the aligned 9O5O cryo-EM structure and used as a reference to calculate the RMSD to the docking results.

One additional docking run was conducted using a single frame extracted from the MD corresponding to 2 ns. In this calculation, Trp216 was set to “Rotate” by applying the default GOLD rotamer library. Every other option was set as described for the ensemble docking calculation.

### 4.2 Docking and MD Simulations Results Analysis

Docking results and frames extracted from the MD simulations were inspected using ChimeraX v1.11^[56,57]^. RMSD of the binding modes obtained with docking was calculated using GOLD, providing the aligned conformation of Compound 4 extracted from the Compound 4-bound K_Ca_2.2 structure as reference. Ligand-protein interactions connected to the binding mode obtained with docking were calculated using PLIP^[58]^.

For the following analysis of the MD simulations, trajectories were aligned on K_Ca_3.1 or K_Ca_2.2 channel Cα atoms (subunits A-D). The VMD v2.0.0a7^[59]^ software was used for visual inspection of the MD trajectories. Root mean square deviation (RMSD), Radius of Gyration (RoG), and interatomic distances during MD simulation were analysed using in-house python scripts based on MDAnalysis^[60]^. Non-covalent interactions during the trajectories were tracked with custom python scripts based on ProLIF^[61]^. *χ*_1_ and *χ*_2_ dihedral angles of Trp322 or Trp216 residues were calculated using the gmx chi function included in GROMACS, after extracting one frame for each nanosecond of the trajectories.

Volume of the S5-Phelix-S6 binding pocket in system K_Ca_3.1_open_P245S was analysed using fpocket^[62]^, a pocket calculation algorithm based on Voronoi tessellation and alpha spheres. First, the trajectory was aligned onto K_Ca_3.1 chain B, where the target pocket is located, and one frame for every nanosecond was extracted. Then, system-wide analysis of the pockets during the MD simulation was conducted. From the resulting density grid the S5-Phelix-S6 pocket location was obtained, using an iso-value of 1.39 (corresponding to the minimum value at which the pocket is not merged with surrounding ones). Finally, the S5-Phelix-S6 pocket location was used to calculate pocket specific descriptors for every frame of the MD simulation.

## ACKNOWLEDGEMENTS

This work was supported by the Research Training Group “Chemical biology of ion channels (Chembion)” funded by the Deutsche Forschungsgemeinschaft (DFG), which is gratefully acknowledged. O.K. was furthermore funded through a Heisenberg-Professorship (DFG; KO 4689/5–2). We gratefully acknowledge the scientific support and HPC resources provided by the Erlangen National High Performance Computing Centre (NHR@FAU) of the Friedrich-Alexander-Universität Erlangen-Nürnberg (FAU) under the NHR project k101ee NHR funding is provided by federal and Bavarian state authorities.

## DATA AVAILABILITY STATEMENT

The video of the Trp216 conformational change in the K_Ca_3.1_open_P245S MD simulation, and the results of the Compund 4 docking calculations are available from the University of Münster datastore: DOI: 10.17879/htsyf-scc57.

## REFERENCES

[1] Gamper, N. e Wang, K. (a c. di) (2021). Pharmacology of Potassium Channels, Springer International Publishing, Cham.

[2] Brown, B.M. et al. (2020). Pharmacology of Small-and Intermediate-Conductance Calcium-Activated Potassium Channels. Annual Review of Pharmacology and Toxicology. 10.1146/annurev-pharmtox-010919-023420.

[3] Kaczmarek, L.K. et al. (2017). International Union of Basic and Clinical Pharmacology. C. Nomenclature and Properties of Calcium-Activated and Sodium-Activated Potassium Channels. Pharmacological Reviews. 10.1124/pr.116.012864.

[4] Fanger, C.M. et al. (1999). Calmodulin Mediates Calcium-dependent Activation of the Intermediate Conductance KCa Channel,IKCa1 *. Journal of Biological Chemistry. 10.1074/jbc.274.9.5746.

[5] Lee, C.-H. e MacKinnon, R. (2018). Activation mechanism of a human SK-calmodulin channel complex elucidated by cryo-EM structures. Science. 10.1126/science.aas9466.

[6] Cassell, S.J. et al. (2025). Mechanism of SK2 channel gating and its modulation by the bee toxin apamin and small molecules. eLife. 10.7554/eLife.107733.

[7] Jedele, S. et al. (2025). Investigating the Dynamics of the KCNN4 Channel: From the Determination of the Complete K+ Permeation Pathway Across the Channel to Its Opening by PIP2. Journal of Chemical Information and Modeling. 10.1021/acs.jcim.4c01711.

[8] Simó-Vicens, R. et al. (2017). A new negative allosteric modulator, AP14145, for the study of small conductance calcium-activated potassium (KCa2) channels. British Journal of Pharmacology. 10.1111/bph.14043.

[9] Lam, J. et al. (2013). The therapeutic potential of small-conductance KCa2 channels in neurodegenerative and psychiatric diseases. Expert Opinion on Therapeutic Targets. 10.1517/14728222.2013.823161.

[10] Ikonen, S. e Riekkinen, P. (1999). Effects of apamin on memory processing of hippocampal-lesioned mice. European Journal of Pharmacology. 10.1016/S0014-2999(99)00616-0.

[11] Holst, A.G. et al. (2024). Inhibition of the KCa2 potassium channel in atrial fibrillation: a randomized phase 2 trial. Nature Medicine. 10.1038/s41591-023-02679-9.

[12] Rahman, M.A. et al. (2023). KCa2.2 (KCNN2): A physiologically and therapeutically important potassium channel. Journal of Neuroscience Research. 10.1002/jnr.25233.

[13] Mulholland, P.J. et al. (2011). Small Conductance Calcium-Activated Potassium Type 2 Channels Regulate Alcohol-Associated Plasticity of Glutamatergic Synapses. Biological Psychiatry. 10.1016/j.biopsych.2010.09.025.

[14] Allen, D. et al. (2011). SK2 Channels Are Neuroprotective for Ischemia-Induced Neuronal Cell Death. Journal of Cerebral Blood Flow & Metabolism. 10.1038/jcbfm.2011.90.

[15] Ataga, K.I. et al. (2021). Haemoglobin response to senicapoc in patients with sickle cell disease: a re-analysis of the Phase III trial. British Journal of Haematology. 10.1111/bjh.17345.

[16] Ataga, K.I. et al. (2011). Improvements in haemolysis and indicators of erythrocyte survival do not correlate with acute vaso-occlusive crises in patients with sickle cell disease: a phase III randomized, placebo-controlled, double-blind study of the gardos channel blocker senicapoc (ICA-17043). British Journal of Haematology. 10.1111/j.1365-2141.2010.08520.x.

[17] Biossil Inc. (2025). A Multicenter, Randomized, Double-blind, Placebo-controlled Study to Determine Efficacy and Safety of SIL-8301 in Sickle Cell Disease (SCD) Patients With a Predominantly Hemolytic Phenotype.

[18] Thi Hong Van, N. e Hyun Nam, J. (2024). Intermediate conductance calcium-activated potassium channel (KCa3.1) in cancer: Emerging roles and therapeutic potentials. Biochemical Pharmacology. 10.1016/j.bcp.2024.116573.

[19] Todesca, L.M. et al. (2024). Targeting KCa3.1 channels to overcome erlotinib resistance in non-small cell lung cancer cells. Cell Death Discovery. 10.1038/s41420-023-01776-5.

[20] Aarhus University Hospital (2026). Phase 0/1 Randomized Clinical Trial of SENIcapoc and PERAmpanel Mono-and Combination Therapy of Newly Diagnosed Glioblastoma.

[21] Sankaranarayanan, A. et al. (2009). Naphtho[1,2-d]thiazol-2-ylamine (SKA-31), a New Activator of KCa2 and KCa3.1 Potassium Channels, Potentiates the Endothelium-Derived Hyperpolarizing Factor Response and Lowers Blood Pressure. Molecular Pharmacology. 10.1124/mol.108.051425.

[22] Coleman, N. et al. (2014). New Positive Ca2+-Activated K+ Channel Gating Modulators with Selectivity for KCa3.1. Molecular Pharmacology. 10.1124/mol.114.093286.

[23] Kolski-Andreaco, A. et al. (2025). VX-445 (elexacaftor) inhibits chloride secretion across human bronchial epithelial cells by directly blocking KCa3.1 channels. PNAS Nexus. 10.1093/pnasnexus/pgaf211.

[24] Devor, D.C. et al. (2022). KCa3.1 potentiation stimulates Cl− secretion in F508del and G551D CFTR-corrected primary human bronchial epithelial cells. American Journal of Physiology-Cell Physiology. 10.1152/ajpcell.00319.2022.

[25] Wulff, H. et al. (2022). Can KCa3.1 channel activators serve as novel inhibitors of platelet aggregation? Journal of Thrombosis and Haemostasis. 10.1111/jth.15863.

[26] Mourre, C. et al. (1997). Apamin, a blocker of the calcium-activated potassium channel, induces neurodegeneration of Purkinje cells exclusively. Brain Research. 10.1016/S0006-8993(97)01165-7.

[27] Nam, Y.-W. et al. (2026). Structural basis for the subtype-selectivity of KCa2.2 channel activators. Nature Communications. 10.1038/s41467-025-67232-3.

[28] Ramanishka, A. et al. (2026). BPS2026 – Structural basis for the subtype-selective activation of KCa3.1 channels. Biophysical Journal. 10.1016/j.bpj.2025.11.2173.

[29] Nguyen, H.M. et al. (2017). Structural Insights into the Atomistic Mechanisms of Action of Small Molecule Inhibitors Targeting the KCa3.1 Channel Pore. Molecular Pharmacology. 10.1124/mol.116.108068.

[30] Henikoff, S. e Henikoff, J.G. (1992). Amino acid substitution matrices from protein blocks. Proceedings of the National Academy of Sciences of the United States of America. 10.1073/pnas.89.22.10915.

[31] Garneau, L. et al. (2014). Aromatic–aromatic interactions between residues in KCa3.1 pore helix and S5 transmembrane segment control the channel gating process. Journal of General Physiology. 10.1085/jgp.201311097.

[32] Hirschberg, B. et al. (1998). Gating of Recombinant Small-Conductance Ca-activated K+ Channels by Calcium. The Journal of General Physiology. 10.1085/jgp.111.4.565.

[33] Madeira, F. et al. (2024). The EMBL-EBI Job Dispatcher sequence analysis tools framework in 2024. Nucleic acids research. 10.1093/nar/gkae241.

[34] Gómez-Tamayo, J.C. et al. (2016). Analysis of the interactions of sulfur-containing amino acids in membrane proteins. Protein Science. 10.1002/pro.2955.

[35] Berman, H.M. et al. (2000). The Protein Data Bank. Nucleic Acids Research. 10.1093/nar/28.1.235.

[36] Vilar, S. et al. (2008). Medicinal Chemistry and the Molecular Operating Environment (MOE): Application of QSAR and Molecular Docking to Drug Discovery. Current Topics in Medicinal Chemistry. 10.2174/156802608786786624.

[37] Halgren, T.A. (1996). Merck molecular force field. I. Basis, form, scope, parameterization, and performance of MMFF94. Journal of Computational Chemistry. 10.1002/(SICI)1096-987X(199604)17:5/6%253C490::AID-JCC1%253E3.0.CO;2-P.

[38] Wu, E.L. et al. (2014). CHARMM-GUI membrane builder toward realistic biological membrane simulations. 10.1002/jcc.23702.

[39] Lee, J. et al. (2016). CHARMM-GUI input generator for NAMD, GROMACS, AMBER, OpenMM, and CHARMM/OpenMM simulations using the CHARMM36 additive force field. Journal of Chemical Theory and Computation. 10.1021/acs.jctc.5b00935.

[40] Lomize, M.A. et al. (2012). OPM database and PPM web server: resources for positioning of proteins in membranes. Nucleic Acids Research. 10.1093/NAR/GKR703.

[41] Jorgensen, W.L. et al. (1983). Comparison of simple potential functions for simulating liquid water. The Journal of Chemical Physics. 10.1063/1.445869.

[42] Abraham, M.J. et al. (2015). GROMACS: High performance molecular simulations through multi-level parallelism from laptops to supercomputers. SoftwareX. 10.1016/J.SOFTX.2015.06.001.

[43] Maier, J.A. et al. (2015). ff14SB: Improving the accuracy of protein side chain and backbone parameters from ff99SB. Journal of Chemical Theory and Computation. 10.1021/acs.jctc.5b00255.

[44] Dickson, C.J. et al. (2022). Lipid21: Complex Lipid Membrane Simulations with AMBER. Journal of Chemical Theory and Computation. 10.1021/acs.jctc.1c01217.

[45] Jakalian, A. et al. (2002). Fast, efficient generation of high-quality atomic charges. AM1-BCC model: II. Parameterization and validation. Journal of Computational Chemistry. 10.1002/jcc.10128.

[46] Wang, J. et al. (2004). Development and testing of a general amber force field. Journal of Computational Chemistry. 10.1002/jcc.20035.

[47] Huang, J. et al. (2017). CHARMM36m: an improved force field for folded and intrinsically disordered proteins. Nature Methods. 10.1038/nmeth.4067.

[48] Vanommeslaeghe, K. et al. (2010). CHARMM General Force Field (CGenFF): A force field for drug-like molecules compatible with the CHARMM all-atom additive biological force fields. Journal of computational chemistry. 10.1002/jcc.21367.

[49] Darden, T. et al. (1993). Particle mesh Ewald: An N·log(N) method for Ewald sums in large systems. The Journal of Chemical Physics. 10.1063/1.464397.

[50] Hess, B. et al. (1997). LINCS: A linear constraint solver for molecular simulations. Journal of Computational Chemistry.

[51] Pedregosa, F. et al. (2011). Scikit-learn: Machine Learning in Python. Journal of Machine Learning Research.

[52] Jones, G. et al. (1997). Development and validation of a genetic algorithm for flexible docking. Journal of Molecular Biology. 10.1006/jmbi.1996.0897.

[53] Verdonk, M.L. et al. (2003). Improved protein–ligand docking using GOLD. Proteins: Structure, Function, and Bioinformatics. 10.1002/prot.10465.

[54] Korb, O. et al. (2009). Empirical Scoring Functions for Advanced Protein−Ligand Docking with PLANTS. Journal of Chemical Information and Modeling. 10.1021/ci800298z.

[55] Jones, G. et al. (1995). Molecular recognition of receptor sites using a genetic algorithm with a description of desolvation. Journal of Molecular Biology. 10.1016/S0022-2836(95)80037-9.

[56] Goddard, T.D. et al. (2018). UCSF ChimeraX: Meeting modern challenges in visualization and analysis. Protein Science. 10.1002/PRO.3235.

[57] Pettersen, E.F. et al. (2021). UCSF ChimeraX: Structure visualization for researchers, educators, and developers. Protein Science. 10.1002/PRO.3943.

[58] Salentin, S. et al. (2015). PLIP: fully automated protein–ligand interaction profiler. Nucleic Acids Research. 10.1093/nar/gkv315.

[59] Humphrey, W. et al. (1996). VMD: Visual molecular dynamics. Journal of Molecular Graphics. 10.1016/0263-7855(96)00018-5.

[60] Gowers, R.J. et al. (2016). MDAnalysis: a python package for the rapid analysis of molecular dynamics simulations. PROC. OF THE 15th PYTHON IN SCIENCE CONF. 10.25080/Majora-629e541a-00e.

[61] Bouysset, C. e Fiorucci, S. (2021). ProLIF: a library to encode molecular interactions as fingerprints. Journal of Cheminformatics. 10.1186/s13321-021-00548-6.

[62] Le Guilloux, V. et al. (2009). Fpocket: An open source platform for ligand pocket detection. BMC Bioinformatics. 10.1186/1471-2105-10-168.

