## Supporting Information for "Mutation-Induced Pocket Deactivation: How Ser353/Pro245 Alters K_Ca_2.2 vs K_Ca_3.1 Ligand Selectivity"

[#] Shared First Authors

* Corresponding author

**Table of Contents**


| | | |
|---|---|---|
| **Table S1** | Available $K_{Ca}2.2$ experimental structures | 3 |
| **Table S2** | Available $K_{Ca}3.1$ experimental structures | 4 |
| **Figure S1** | Sequence alignment of $K_{Ca}2.2$ and $K_{Ca}3.1$ channels across different species | 5 |
| **Table S3** | Average RMSD values obtained in the $K_{Ca}3.1$ MD simulations | 6 |
| **Figure S2** | $K_{Ca}3.1$ channel subunits RMSD values throughout the MD simulations | 7 |
| **Figure S3** | Calmodulin subunits RMSD variation throughout the MD simulations | 8 |
| **Figure S4** | $K_{Ca}3.1$ channel Radius of Gyration values throughout the MD simulations | 9 |
| **Figure S5** | Plots of the $\chi_1$ and $\chi_2$ dihedral angles of residue Trp322 during the $K_{Ca}2.2$ MD simulations | 10 |
| **Figure S6** | Trp216-Ser245 distances throughout the KCa3.1_open_P245S MD simulations | 11 |
| **Figure S7** | S5-Phelix-S6 Pocket calculated from the $K_{Ca}2.2$ and $K_{Ca}3.1$ channels | 12 |
| **Figure S8** | Results of the docking calculation of Compound 4 to the frame corresponding to the Trp216 *conformation a* | 13 |
| **Table S4** | Details of Molecular Dynamics systems | 14 |

**Table S1**: $K_{Ca}2.2$ structures published on the RCSB PDB database (https://www.rcsb.org/, accessed on 21.03.2026). For each structure the following details are indicated: PDB ID; species from which the $K_{Ca}2.2$ and calmodulin sequences were obtained; hydrophobic gate state (open/closed, based on distance between residue V390 in opposite subunits); co-determined ions; co-determine ligands (lipid molecules are excluded from this field); Trp322 orientation (defined as "Upward" if the side chain is directed towards the extracellular side, and "Downward" if the side chain is directed towards the intracellular side); mutations in the sequence; DOI of the reference paper.
* Ca2+ ions are bound to calmodulin C-Lobe but not to the N-Lobe.

| PDB ID | Species | Determination Techniques | Resolution (Å) | Hydrophobic Gate State | Co-determined Ions | Co-determined Ligands | Trp322 Orientation | Sequence Mutation | Reference DOI |
|---|---|---|---|---|---|---|---|---|---|
| 8V2G | Rattus norvegicus | Cryo-EM | 3.18 | Open | $K^+$, $Ca^{2+}$ | / | Upward | / | 10.1038/s41467-025-59061-1 |
| 8V2H | Rattus norvegicus | Cryo-EM | 3.1 | Closed | $K^+$, $Ca^{2+}$ | AP14145 | Upward | / | 10.1038/s41467-025-59061-1 |
| 8V3G | Rattus norvegicus | Cryo-EM | 3.1 | Open | $K^+$, $Ca^{2+}$ | UCL1684 | Upward | / | 10.1038/s41467-025-59061-1 |
| 9EIO | Rattus norvegicus | Cryo-EM | 3.62 | Open | $K^*$, $Ca^{2+}$ | / | Downward | Mutation F244S | 10.1038/s41467-025-59061-1 |
| 9O48 | Homo sapiens | Cryo-EM | 3.1 | Open | $K^+$, $Ca^{2+}$ | / | Upward | / | 10.7554/eLife.107733 |
| 9O51 | Homo sapiens | Cryo-EM | 3.4 | Closed | $K^+$, $Ca^{2+}$ * | / | Upward | / | 10.7554/eLife.107733 |
| 9O52 | Homo sapiens | Cryo-EM | 3.18 | Closed | $K^+$, $Ca^{2+}$ | Apamin | Upward | / | 10.7554/eLife.107733 |
| 9O53 | Homo sapiens | Cryo-EM | 3.3 | Closed | $K^+$, $Ca^{2+}$ | Compound 1 | Upward | / | 10.7554/eLife.107733 |
| 9O5O | Homo sapiens | Cryo-EM | 3.1 | Open | $K^+$, $Ca^{2+}$ | Compound 4 | Upward | / | 10.7554/eLife.107733 |
| 9O7S | Rattus norvegicus | Cryo-EM | 2.71 | Open | $K^+$, $Ca^{2+}$ | NS309 | Upward | / | 10.1038/s41467-025-67232-3 |
| 9O85 | Rattus norvegicus | Cryo-EM | 3.13 | Open | $K^+$, $Ca^{2+}$ | Rimtuzalcap | Upward | / | 10.1038/s41467-025-67232-3 |
| 9O93 | Rattus norvegicus | Cryo-EM | 2.96 | Closed | $K^+$ | Rimtuzalcap | Upward, flipped | / | 10.1038/s41467-025-67232-3 |
| 9VU9 | Homo sapiens | Cryo-EM | 3.34 | Closed | $K^+$ | / | Upward | / | 10.1038/s41467-026-68475-4 |
| 9VUA | Homo sapiens | Cryo-EM | 3.23 | Closed | $K^+$ | AP30663 | Upward | / | 10.1038/s41467-026-68475-4 |
| 9VUB | Homo sapiens | Cryo-EM | 3.35 | Open | $K^+$, $Ca^{2+}$ | Rimtuzalcap | Upward | / | 10.1038/s41467-026-68475-4 |
| 9VUC | Homo sapiens | Cryo-EM | 2.96 | Closed | $K^+$ | UCL1684 | Upward | | 10.1038/s41467-026-68475-4 |

**Table S2:** $K_{Ca}3.1$ structures published on the RCSB PDB database (https://www.rcsb.org/, accessed on 21.03.2026). For each structure the following details are indicated: PDB ID; species from which the $K_{Ca}3.1$ and calmodulin sequences were obtained; hydrophobic gate state (open/closed, based on distance between residue V282 in opposite subunits); co-determined ions; co-determine ligands (lipid molecules are excluded from this field); Trp216 orientation (defined as "Upward" if the side chain is directed towards the extracellular side, and "Downward" if the side chain is directed towards the intracellular side); mutations in the sequence; DOI of the reference paper.

| PDB ID | Species | Determination Techniques | Resolution (Å) | Hydrophobic Gate State | Co-determined Ions | Co-determined Ligands | Trp216 Orientation | Sequence Mutation | Reference DOI |
|---|---|---|---|---|---|---|---|---|---|
| 6CNM | Homo sapiens | Cryo-EM | 3.4 | Closed | $K^+$ | / | Downward | / | 10.1126/science.aas9466 |
| 6CNN | Homo sapiens | Cryo-EM | 3.5 | Closed | $K^+$, $Ca^{2+}$ | / | Downward | / | 10.1126/science.aas9466 |
| 6CNO | Homo sapiens | Cryo-EM | 4.7 | Open | $Ca^{2+}$ | / | Downward | / | 10.1126/science.aas9466 |
| 9ED1 | Homo sapiens, Rattus norvegicus | Cryo-EM | 3.5 | Closed | $K^+$, $Ca^{2+}$ | DHP-103 | Downward | / | 10.1073/pnas.2425494122 |
| 9OA8 | Homo sapiens, Rattus norvegicus | Cryo-EM | 3.59 | Open | $K^+$, $Ca^{2+}$ | NS309 | Downward | / | 10.1038/s41467-025-67232-3 |
| 9YDZ | Homo sapiens | Cryo-EM | 3.4 | Closed | $K^+$ | Rimtuzalcap | Downward | Mutation R355K | 10.1038/s41467-025-67232-3 |
| 9Y5Q | Homo sapiens | Cryo-EM | 4.73 | Open | $K^+$, $Ca^{2+}$ | Rimtuzalcap | Downward | Mutation R355K | 10.1038/s41467-025-67232-3 |

**Figure S1:** Multi sequence alignment (MSA) of $K_{Ca}$2.2 and $K_{Ca}$3.1 channel sequences across different species. Residues corresponding to Ser353 in the human $K_{Ca}$2.2 and Pro245 in the human $K_{Ca}$3.1 are highlighted. Sequences were retrieved from the Uniprot database (https://www.uniprot.org/), and correspond to the following Uniprot codes (same order as in the figure): Q9H2S1, P58390, P70604, A0A6I8RA55, A0AC58GLV1, A0A8M2B2K1, A0AB32U3U6, O15554, O89109, Q9QYW1, A0A6I8QL32, X1WCL6. The sequence alignment was computed using the EMBL-EBI Clustal Omega web server (https://www.ebi.ac.uk/jdispatcher/msa/clustalo).

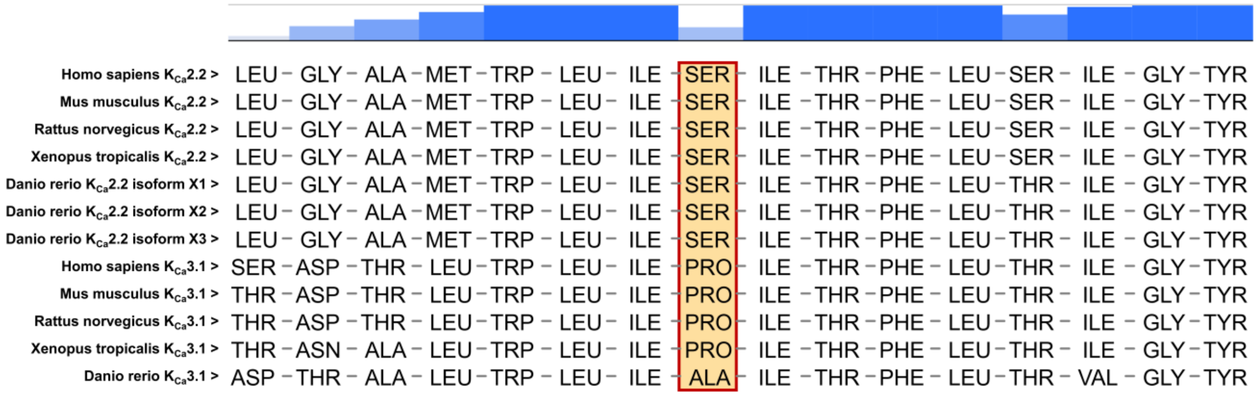

**Table S3:** Average RMSD values, and relative standard deviation, of the $K_{Ca}3.1$ channel subunits and the calmodulin subunits across the MD simulations. Trajectories were aligned on the Cα atoms of the $K_{Ca}3.1$ channel subunits, using the first frame of the production phase as a reference.

| | | **RMSD ± sd (Å)** | |
|---|---|---|---|
| **MD System** | **MD replica** | **$K_{Ca}3.1$ Channel Subunits** | **Calmodulin Subunits** |
| $K_{Ca}3.1$_closed_WT | Run 1 | 3.804 ± 0.402 | 5.315 ± 0.877 |
| | Run 2 | 3.682 ± 0.213 | 5.254 ± 0.771 |
| | Run 3 | 3.718 ± 0.330 | 5.652 ± 1.036 |
| $K_{Ca}3.1$_closed_P245S | Run 1 | 3.749 ± 0.445 | 5.695 ± 0.741 |
| | Run 2 | 3.634 ± 0.393 | 5.438 ± 0.649 |
| | Run 3 | 3.586 ± 0.271 | 5.569 ± 0.660 |
| $K_{Ca}3.1$_open_WT | Run 1 | 4.578 ± 0.490 | 4.469 ± 0.376 |
| | Run 2 | 3.833 ± 0.284 | 4.608 ± 0.393 |
| | Run 3 | 3.805 ± 0.275 | 4.330 ± 0.339 |
| $K_{Ca}3.1$_open_P245S | Run 1 | 4.171 ± 0.409 | 4.304 ± 0.337 |
| | Run 2 | 3.704 ± 0.317 | 4.166 ± 0.348 |
| | Run 3 | 3.961 ± 0.253 | 4.400 ± 0.381 |

**Figure S2:** Root mean square deviation (RMSD) values calculated for the $K_{Ca}3.1$ channel Cα atoms across the triplicate MD simulations of the systems **A**) $K_{Ca}3.1$_closed_WT, **B**) $K_{Ca}3.1$_closed_P245S, **C**) $K_{Ca}3.1$_open_WT, and **D**) $K_{Ca}3.1$_open_P245S. Trajectories were aligned on the Cα atoms of the $K_{Ca}3.1$ channel subunits, using the first frame of the production phase as a reference.

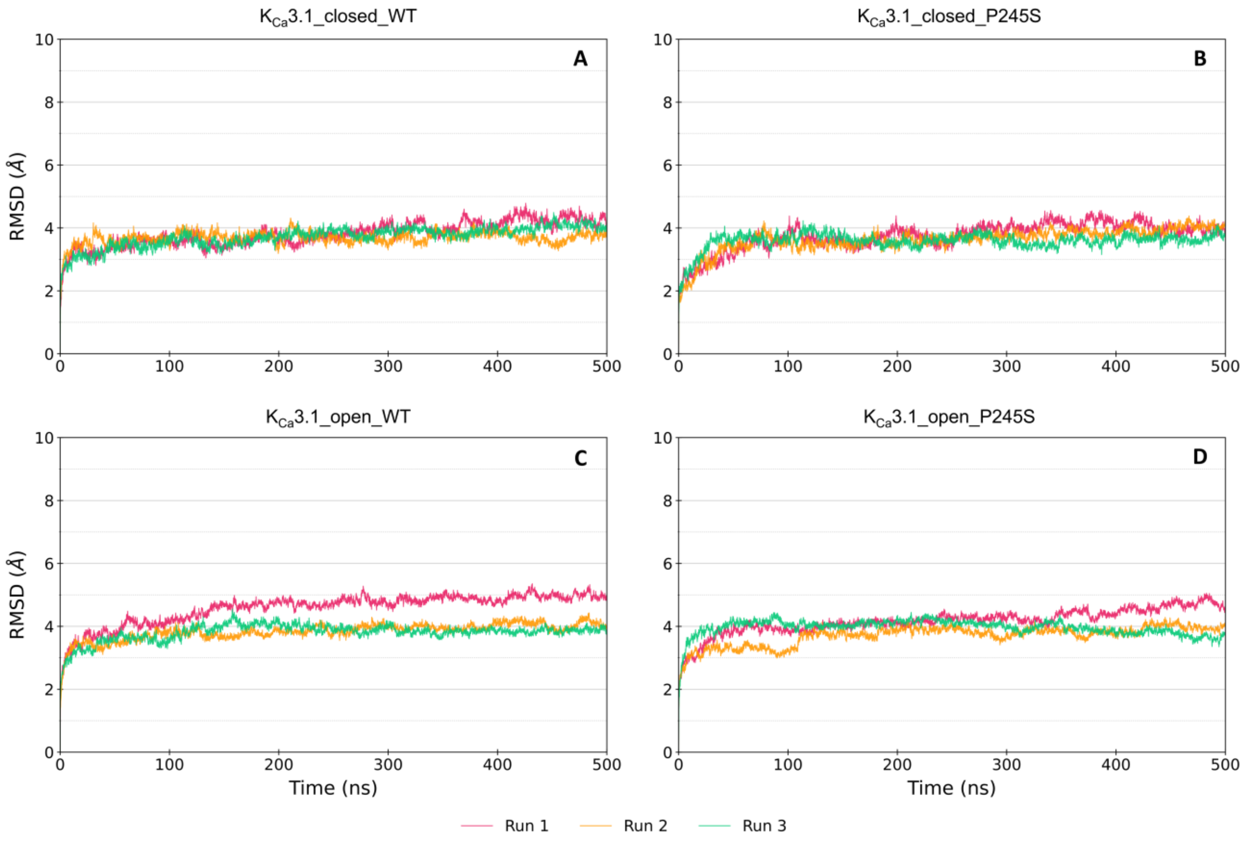

**Figure S3:** Root mean square deviation (RMSD) values calculated for the calmodulin subunits Cα atoms across the triplicate MD simulations of the systems **A**) $K_{Ca}$3.1_closed_WT, **B**) $K_{Ca}$3.1_closed_P245S, **C**) $K_{Ca}$3.1_open_WT, and **D**) $K_{Ca}$3.1_open_P245S. Trajectories were aligned on the Cα atoms of the $K_{Ca}$3.1 channel subunits, using the first frame of the production phase as a reference.

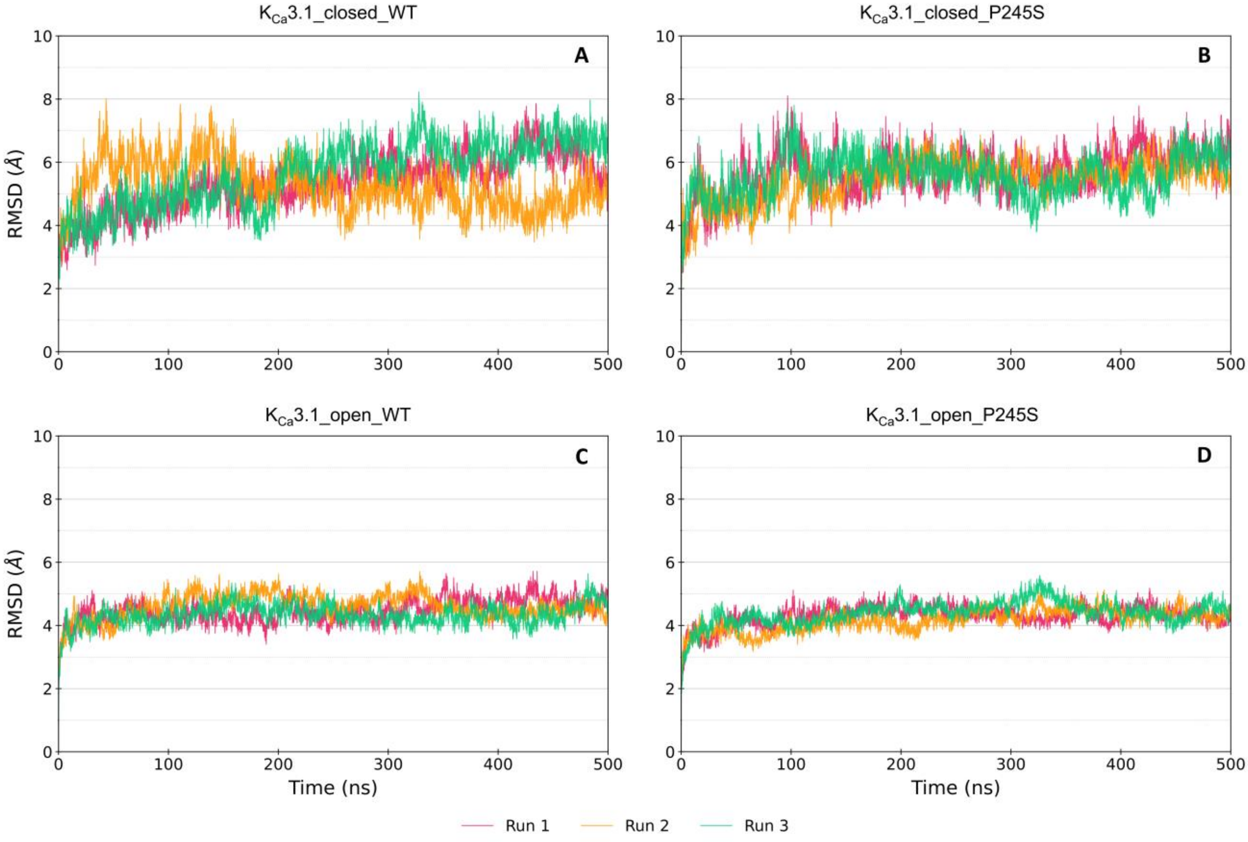

**Figure S4:** Radius of gyration (RoG) values calculated for the $K_{Ca}3.1$ channel subunits across the triplicate MD simulations of the systems **A**) $K_{Ca}3.1$_closed_WT, **B**) $K_{Ca}3.1$_closed_P245S, **C**) $K_{Ca}3.1$_open_WT, and **D**) $K_{Ca}3.1$_open_P245S. Trajectories were aligned on the Cα atoms of the $K_{Ca}3.1$ channel subunits, using the first frame of the production phase as a reference.

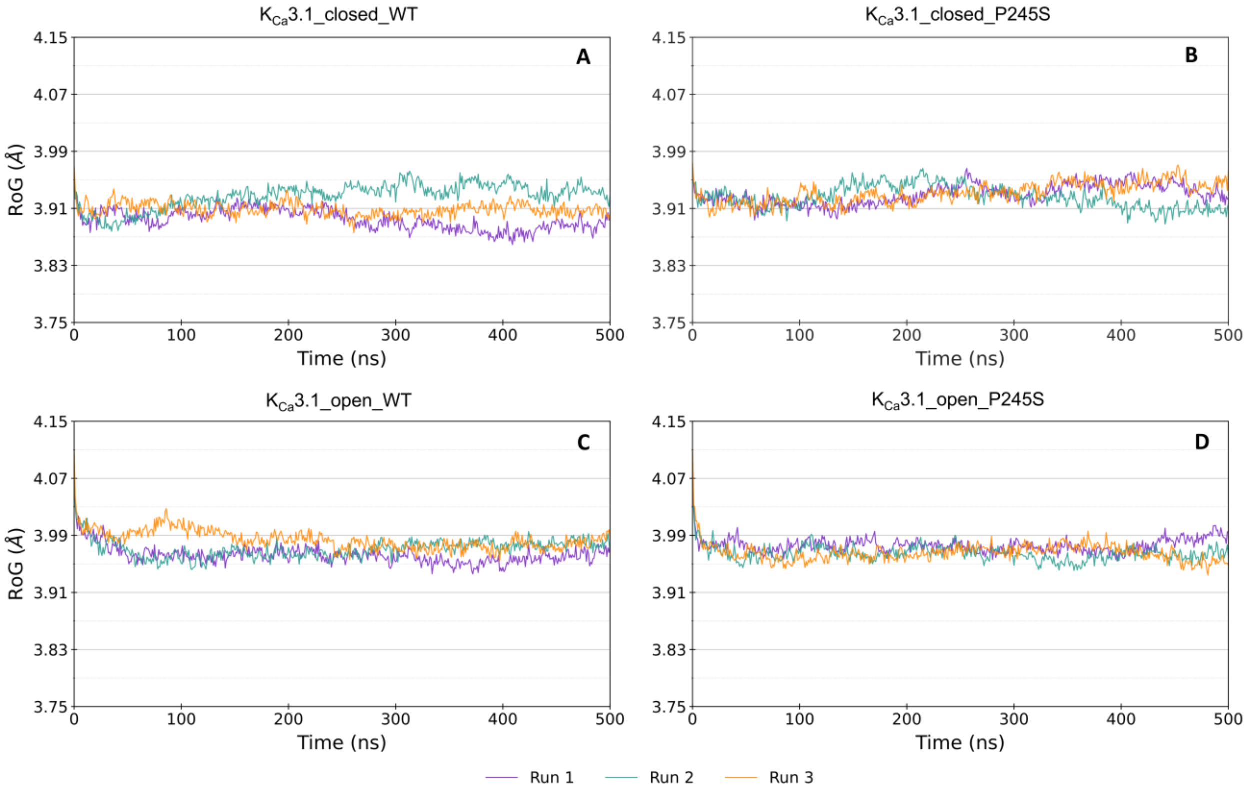

**Figure S5:** Plots of the $\chi_1$ and $\chi_2$ dihedral angles of residue Trp322 during the $K_{Ca}2.2$ MD simulations. Each dot represents a combination of dihedral angles in a specific frame of the trajectory. Time evolution along the simulation is represented by colour shift from yellow to green. Reference dihedral angles obtained from the Compound 4-bound $K_{Ca}2.2$ and $K_{Ca}3.1$_open cryo-EM structures are indicated with a blue and red cross, respectively. **A**) System Compound 1-bound $K_{Ca}2.2$, run 1, chain A. **B**) System Compound 4-bound $K_{Ca}2.2$, run 1, chain A.

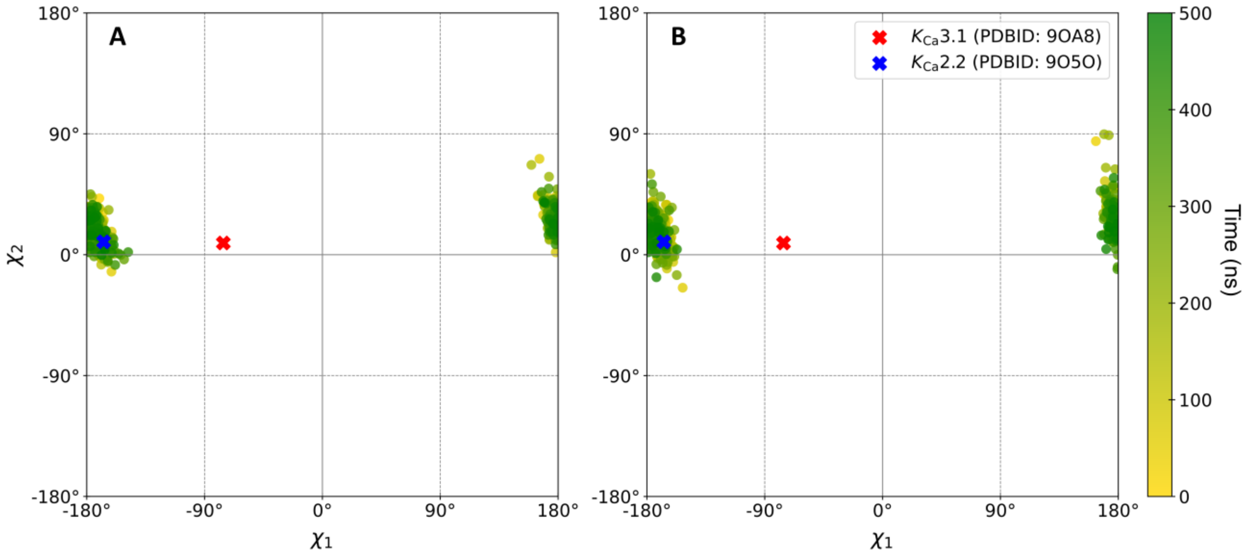

**Figure S6:** Distance between the Trp216 and the Ser245 Cα atoms in the distinct $K_{Ca}3.1$ chains, during the KCa3.1_open_P245S MD simulation, run2.

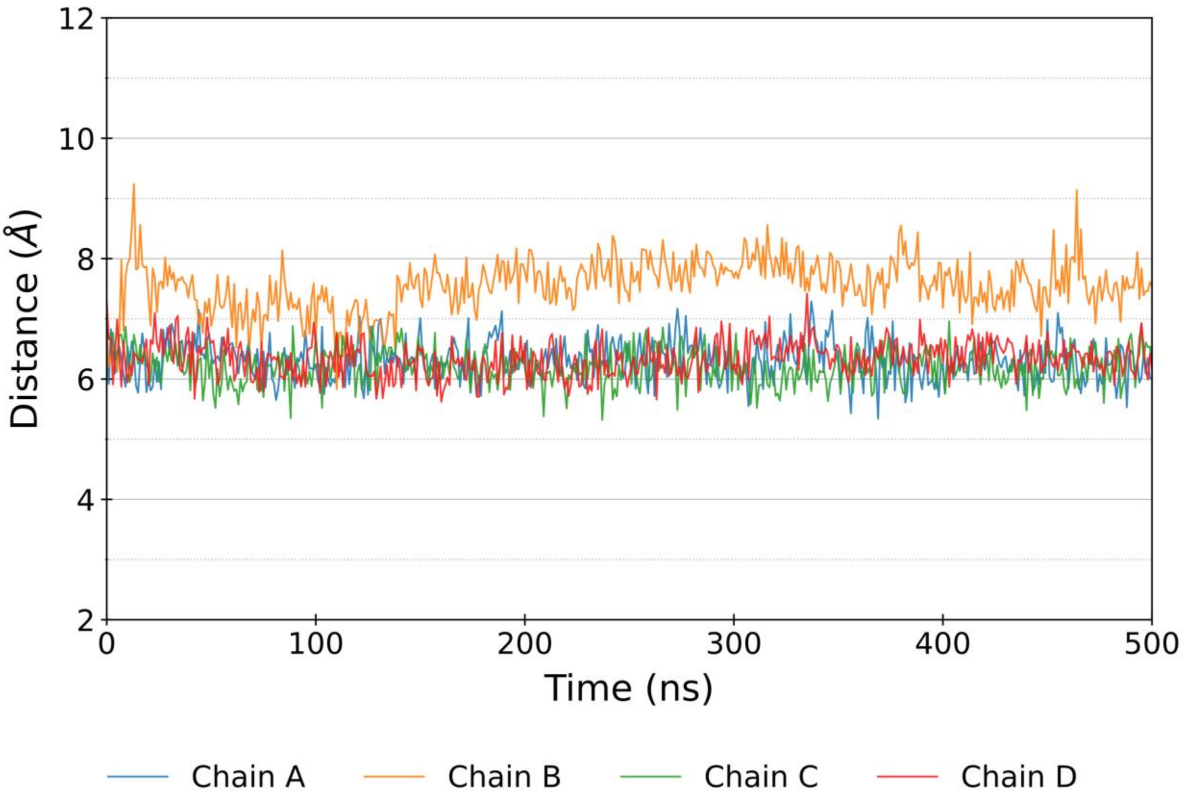

**Figure S7:** Pockets calculated with the fpocket software from the frames extracted from the $K_{Ca}3.1$ MD simulations and from the $K_{Ca}2.2$ structure. The cryo-EM binding mode of Compound 4 (PDB ID: 9O5O) is represented with grey sticks, as a reference. $K_{Ca}3.1$ and $K_{Ca}2.2$ channels are represented with green ribbons and sticks. With blue surfaces are represented the pockets calculated in the structure extracted from the KCa3.1_open_P245S simulation (run2) corresponding to Trp216 **A**) *conformation a*, **B**) *conformation b*, **C**) *conformation c*, and **D**) the pockets calculated from the Compound 4-bound $K_{Ca}2.2$ structure (PDB ID: 9O5O). Only pockets within 5 Å from Compound 4 are included in the figure.

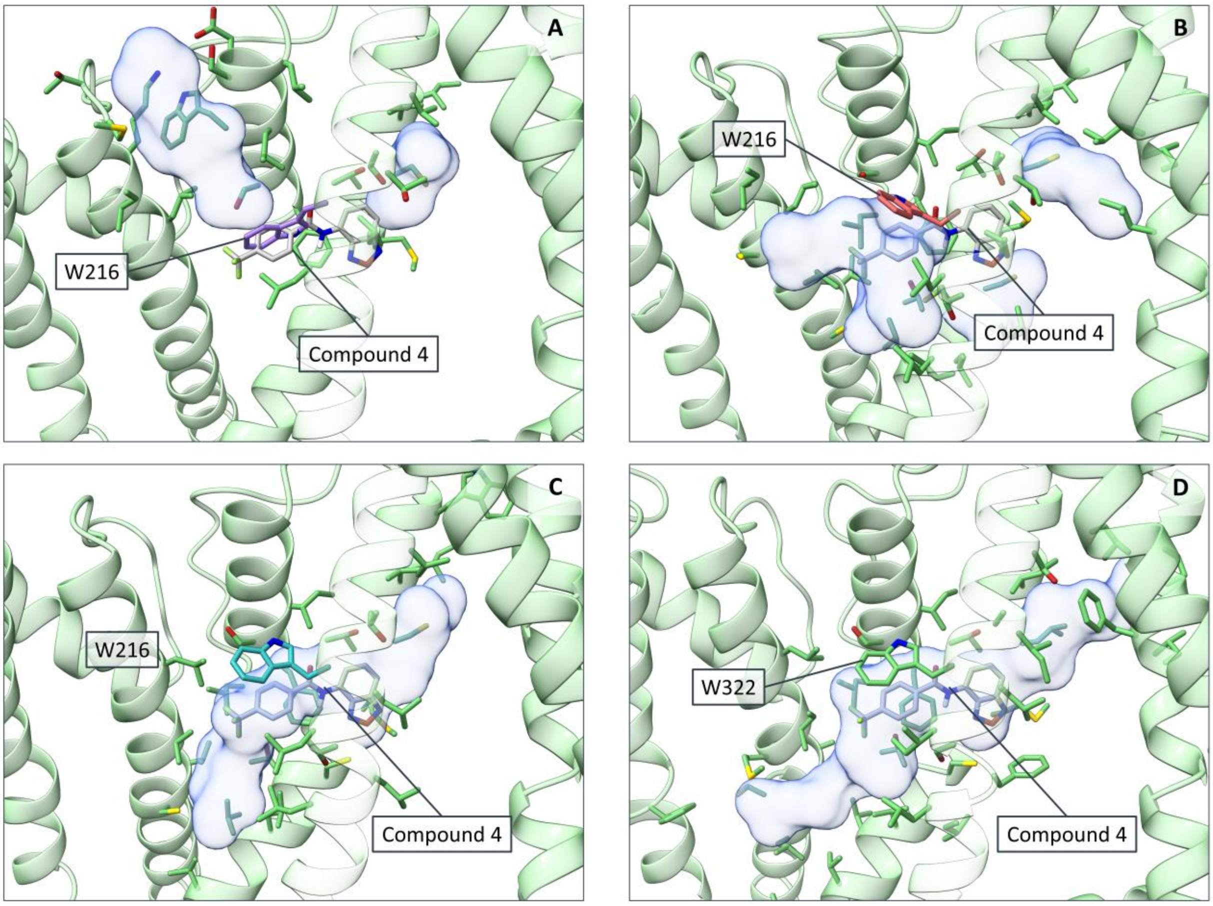

**Figure S8:** Docking poses obtained by docking Compound 4 to the conformation extracted from the $K_{Ca}3.1$_open_P245S MD simulation (run2, chain B) corresponding to 2 ns (Trp216 *conformation a*). $K_{Ca}3.1$ channel is represented with orange ribbons. The reference binding mode of compound 4 extracted from the Compound 4-bound $K_{Ca}2.2$ cryo-EM structure (PDB ID: 9O5O) is represented with grey sticks. Representative conformations of the 100 binding modes obtained through the docking calculation are represented with transparent light-blue sticks.

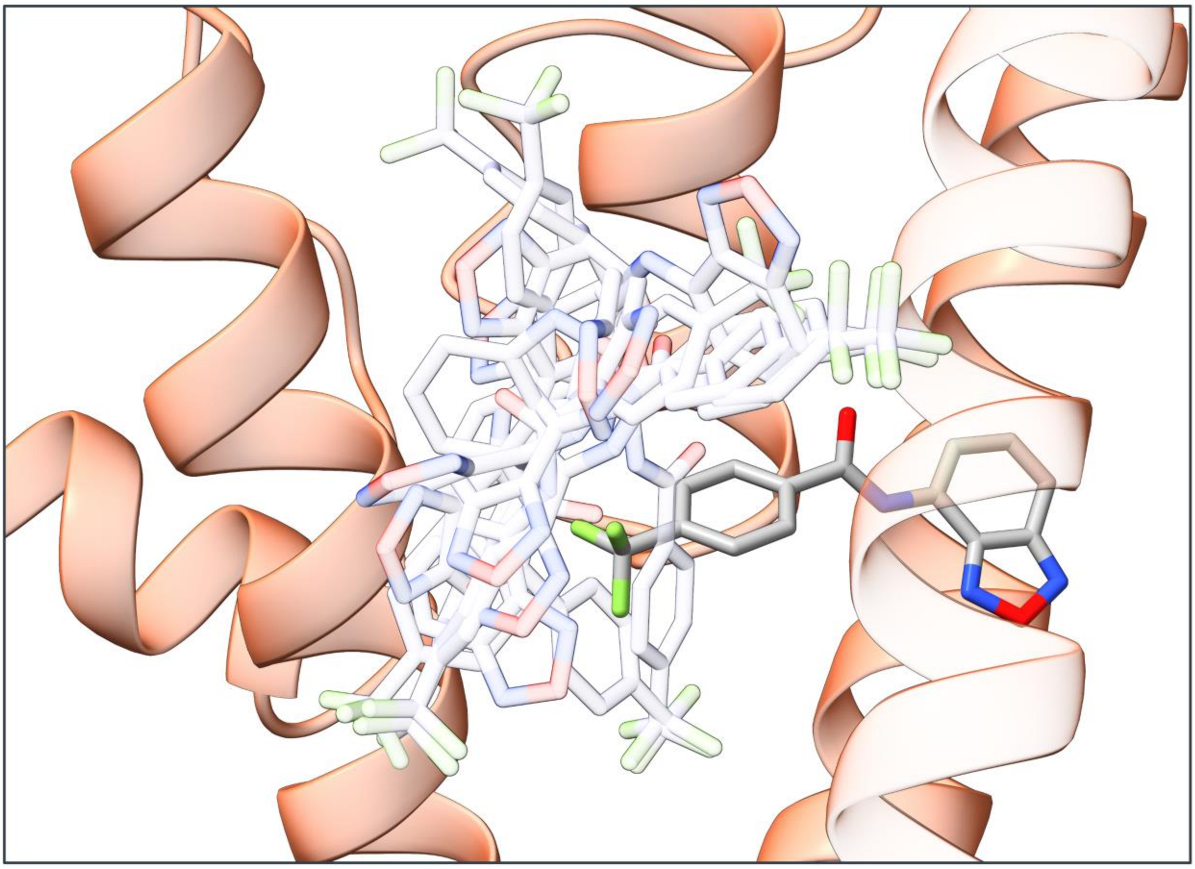

**Table S4:** Details of the MD systems $K_{Ca}3.1$_closed_WT, $K_{Ca}3.1$_closed_P245S, $K_{Ca}3.1$_open_WT, $K_{Ca}3.1$_open_P245S, Compound 1-bound $K_{Ca}2.2$, and Compound 4-bound $K_{Ca}2.2$. For each system, the following details are indicated: system name; initial dimensions of the box along the X, Y, and Z axis; number of POPC molecules contained in each membrane leaflet; number of $K^+$ and $Cl^-$ ions added to the system; total number of water molecules; total number of atoms in each system.

| | Initial System Dimensions (Å) | | | N° POPC | | N° Ions | | | |
|---|---|---|---|---|---|---|---|---|---|
| **System Name** | **X** | **Y** | **Z** | **Outer Leaflet** | **Inner Leaflet** | **$K^+$** | **$Cl^-$** | **Water Molecules** | **Total Atoms** |
| $K_{Ca}3.1$_closed_WT | 160.08 | 160.08 | 150.88 | 314 | 306 | 233 | 274 | 82,293 | 359,005 |
| $K_{Ca}3.1$_closed_P245S | 160.08 | 160.08 | 150.88 | 314 | 306 | 233 | 274 | 82,156 | 358,582 |
| $K_{Ca}3.1$_open_WT | 165.11 | 165.11 | 149.16 | 337 | 327 | 244 | 251 | 84,195 | 375,571 |
| $K_{Ca}3.1$_open_P245S | 165.1 | 165.1 | 149.25 | 337 | 327 | 244 | 251 | 84,278 | 375,808 |
| Compound 1-bound $K_{Ca}2.2$ | 160.11 | 160.11 | 131.89 | 317 | 303 | 189 | 205 | 65,152 | 311,584 |
| Compound 4-bound $K_{Ca}2.2$ | 160.12 | 160.12 | 130.38 | 320 | 302 | 186 | 200 | 64,273 | 309,077 |
